# Time-dynamics of mitochondrial membrane potential reveal an inhibition of ATP synthesis in mitosis

**DOI:** 10.1101/772244

**Authors:** Joon Ho Kang, Georgios Katsikis, Max A. Stockslager, Daniel Lim, Michael B. Yaffe, Scott R. Manalis, Teemu P. Miettinen

## Abstract

The energetic demands of a cell are believed to increase during mitosis ^1–7^. As cells transit from G2 into mitosis, mitochondrial electron transport chain (ETC) activity increases ^4,8,9^, and cellular ATP levels progressively decrease until the metaphase-anaphase transition ^3,7,10^, consistent with elevated consumption. The rates of ATP synthesis during mitosis, however, have not been quantified. Here, we monitor mitochondrial membrane potential of single lymphocytes and demonstrate that cyclin-dependent kinase 1 (CDK1) activity causes mitochondrial hyperpolarization from G2/M until the metaphase-anaphase transition. By using an electrical circuit model of mitochondria, we quantify the time-dynamics of mitochondrial membrane potential under normal and perturbed conditions to extract mitochondrial ATP synthesis rates in mitosis. We found that mitochondrial ATP synthesis decreases by approximately 50 % during early mitosis, when CDK1 is active, and increases back to G2 levels during cytokinesis. Consistently, acute inhibition of mitochondrial ATP synthesis failed to delay cell division. Our results provide a quantitative understanding of mitochondrial bioenergetics in mitosis and challenge the traditional dogma that cell division is a highly energy demanding process.

## Main text

In animal cells, including most cancer cells, energy in the form of ATP is produced mainly through oxidative phosphorylation in mitochondria (reviewed in ^11–13^). However, largely due to a lack of quantitative single-cell approaches, little is known about ATP synthesis during mitosis. To study mitochondrial bioenergetics at the single-cell level, we combined suspended microchannel resonators (SMR), a non-invasive single-cell buoyant mass sensor, with a fluorescence detection system. This allowed us to monitor cell mass-normalized fluorescence signals with a temporal resolution of 2 min and a fluorescence measurement error of 2 % without perturbing normal growth ^14,15^ (Supplementary Fig. 1, Supplementary note 1). Using this setup, we examined the murine lymphocytic leukemia cell line L1210 grown in the presence of non-quenching concentrations (10 nM) of TMRE, a fluorescent probe for mitochondrial membrane potential (ΔΨm) ^16^ (Supplementary Fig. 2A, Supplementary note 2). Monitoring the mass-normalized TMRE signal over multiple cell generations revealed oscillatory TMRE behavior with a robust, transient and extensive spike-like increase in TMRE signal preceding the end of each cell cycle (Fig. 1A). The rate of TMRE increase and decrease were not limited by TMRE diffusion speed (Supplementary Figs. 2C-E, Supplementary note 2). Similar TMRE behavior persisted in both glucose and galactose-based culture media and in other cell types, including mouse BaF3 pro-B lymphocytes, chicken DT40 lymphoblasts, suspension HeLa cells and, importantly, in primary CD8+ and CD3+ human T-cells (Supplementary Fig. 3). However, the mouse Fl5.12 pro-B lymphocytes did not display any change in TMRE at the end of cell cycle (Supplementary Fig. 3A). L1210 cells were utilized as model system in all further studies.

**Fig. 1:**
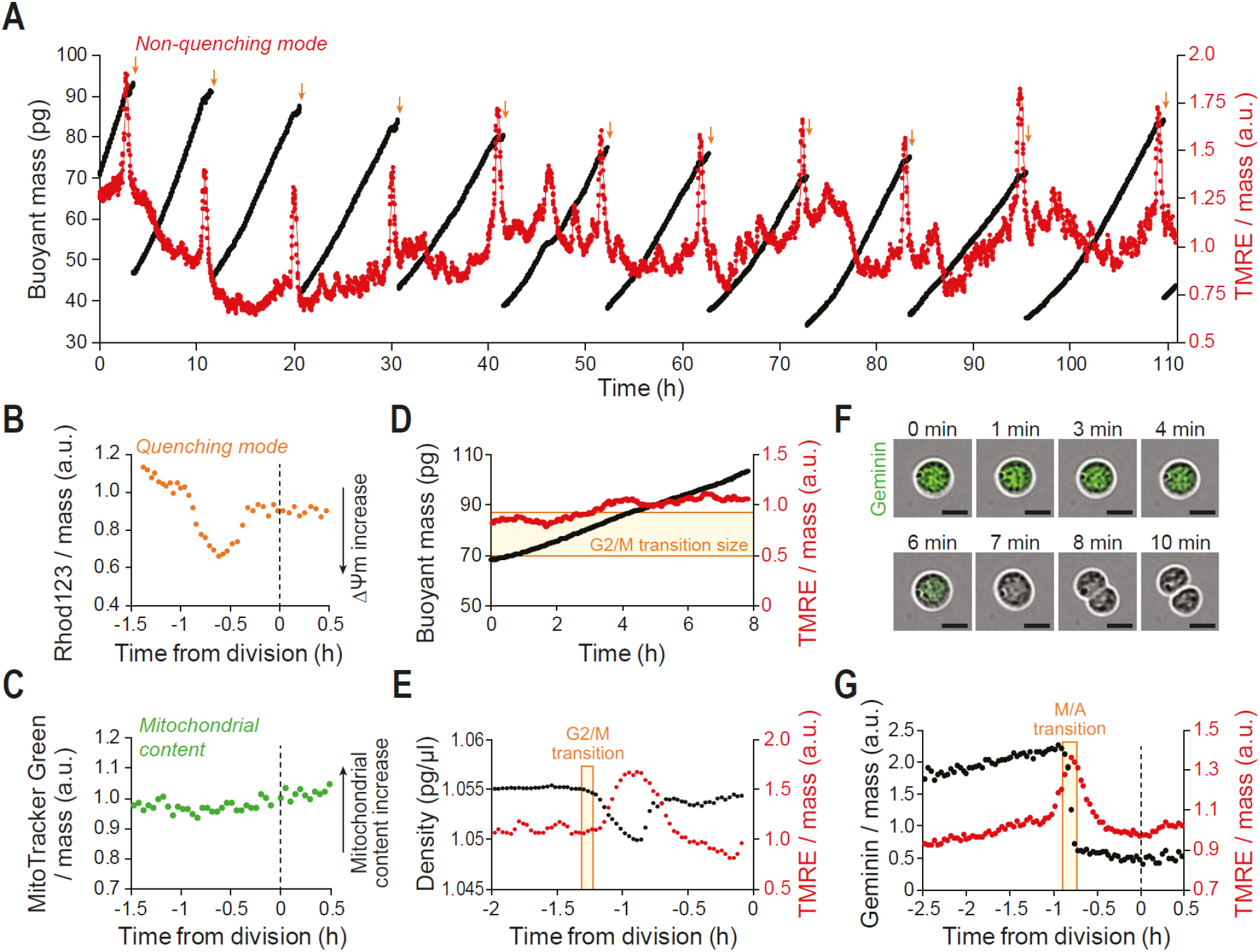
Mitochondria transiently hyperpolarize during prophase and metaphase. **(A)** Buoyant mass (black) and mass-normalized TMRE (red) trace for a single L1210 cell and its progeny over 10 full generations with a measurement interval of 1.9 min. At each cell division (orange arrows), one of the daughter cells is randomly kept and monitored, while the other is discarded. TMRE was used in a non-quenching concentration (10 nM). **(B)** Mass-normalized Rhod132 trace for a L1210 cell around cell division. Cells were loaded with a quenching concentration of Rhod123 (10 μM) and immediately analyzed with no Rhod123 in the culture media. In the quenching mode, the fluorescence signal behavior is reversed. **(C)** Mass-normalized MitoTracker Green (50 nM) trace for a L1210 cell around cell division. **(D)** Buoyant mass (black) and mass-normalized TMRE (red) trace for a L1210 cell treated with 2.5 μM RO-3306 to inhibit mitotic entry. Typical size for mitotic entry is illustrated with light yellow area. **(E)** Cell density (black) and mass-normalized TMRE (red) trace for a L1210 cell around cell division. Mitotic entry (G2/M transition) is illustrated with light yellow area. **(F)** Representative phase contrast (grey) and mAG-hGeminin cell cycle reporter (green) images of a L1210 cell in mitosis. Scale bars denote 10 μm. **(G)** Mass-normalized mAG-hGeminin (black) and mass-normalized TMRE (red) trace for a L1210 cell around cell division. Metaphase-to-anaphase transition (M/A transition) is illustrated with light yellow area.

To validate that the spike-like TMRE increase reflects an increase in ΔΨm, we used quenching concentrations (10 μM) of an alternative ΔΨm probe, Rhod123 (Supplementary Fig. 2B, Supplementary note 2). Rhod123 signal suddenly decreased approximately 1h prior to cell division (Fig. 1B), consistent with a spike-like increase in ΔΨm. In contrast, the ΔΨm-insensitive MitoTracker Green probe did not display any changes at the end of cell cycle (Fig. 1C). We next examined changes in plasma membrane potential (ΔΨp) using a 1 μM DiBAC_4_(3) probe. The DiBAC_4_(3) signal was reduced before and during the TMRE increase, indicative of increased ΔΨp (Supplementary Fig. 4A). We speculated that this ΔΨp change may be a feature of mitotic cell swelling ^17,18^, which can be inhibited with 5-(N-ethyl-N-isopropyl)amiloride (EIPA) ^17^. Indeed, treatment of L1210 cells with 5 μM EIPA partially inhibited the observed decrease in the DiBAC_4_(3) signal but did not affect the TMRE signal increase (Supplementary Fig. 4). This indicates that mitotic cell swelling associates with changes in plasma membrane potential, but the extent to which plasma membrane hyperpolarizes is not affecting the reliability of TMRE as a reporter for ΔΨm (Supplementary note 2).

Next, we studied the exact timing of the ΔΨm increase. Inhibition of mitotic entry using 2.5 μM CDK1 inhibitor RO-3306 completely eliminated the observed increase in TMRE signal, despite the fact that cells continued to increase in size beyond the typical G2/M transition (Fig. 1D). We then monitored single-cell density to compare TMRE signal increase to the timing of mitotic cell swelling, an event that is known to start in prophase ^17,18^. The increase in the TMRE signal was observed immediately following the onset of density reduction, indicating that mitochondrial hyperpolarization begins shortly after mitotic entry (Fig. 1E). This timing of TMRE increase was further validated using biophysical markers of G2/M transition (Supplementary Fig. 5) ^14,15^. We next compared the timing of TMRE signal increase to the degradation of the protein Geminin using L1210 FUCCI cells, which express fluorescently labelled Geminin (Geminin-mAG) ^15,19^. The Geminin-mAG signal was fully degraded in approximately 8.6 minutes at the metaphase-anaphase transition (Fig. 1F, Supplementary Fig. 6), and the loss of Geminin aligned exactly with the maximal TMRE signal (Fig. 1G). In most cells, the TMRE signal declined back to typically G2 levels before the final abscission of the daughter cells (Figs. 1E, 1G). Together, these results indicate that the mitochondrial hyperpolarization begins shortly after the G2/M transition, reaches a maximum at the metaphase-anaphase transition and returns to G2 levels during cytokinesis.

Previous work has shown that the CDK1/Cyclin B complex localizes to mitochondria during mitosis and directly phosphorylates components of the mitochondrial ETC ^4^. CDK1 also associates with other metabolic proteins, including the α subunit of ATP synthase ^20^. CDK1 activity is minimal before mitotic entry, after which the switch-like activation of CDK1/cyclin B complex results in high CDK1 activity until the onset of anaphase ^21–23^. Since the timing of mitochondrial hyperpolarization coincided exactly with the reported CDK1 activity, we hypothesized that the switch-like CDK1 activity was causally responsible for mitochondrial hyperpolarization. To test this, we first arrested cells in a CDK1 active-state (prometaphase and metaphase) using three different chemicals: the kinesin motor inhibitor S-trityl-l-cysteine (STLC), the microtubule polymerization inhibitor nocodazole, and the anaphase-promoting complex inhibitor proTAME. In response to any of these three chemicals we observed that TMRE signal increased following mitotic entry and plateaued to a high level during the mitotic arrest, indicative of mitochondria reaching a steady, hyperpolarized state. Next, we partially inhibited CDK1 with RO-3306 (1 μM) or with an alternative CDK1 inhibitor BMS-265246 (400 nM) and examined the level of TMRE during STLC mediated prometaphase arrest. Note that complete inhibition of CDK1 blocks mitotic entry, but it is possible to partially inhibit CDK1 while allowing mitotic entry and progression ^15,17^. We observed that CDK1 inhibition reduces the mitotic mitochondrial hyperpolarization (Figs. 2B, 2C). Finally, we arrested cells in prometaphase with STLC and after the TMRE signal had reached a new equilibrium in mitosis we treated the cells with 100 nM okadaic acid (O.A.) to inhibit the protein phosphatase PP2A and block the dephosphorylation of CDK1 targets ^23^. The O.A. treatment increased TMRE signal (Fig. 2D). In contrast, when the CDK1 activity of prometaphase-arrested cells was inhibited with 5 μM RO-3306 the TMRE signal returned to G2 levels (Fig. 2E). Together, these results indicate that CDK1 activity drives the mitochondrial hyperpolarization in early mitosis.

CDK1 has been suggested to promote mitochondrial ATP synthesis ^4^. Considering the prevailing dogma that mitosis is energy intensive ^1–7^, we studied whether acute inhibitions of mitochondrial ATP synthesis affected cell division. Direct measurements of oxygen consumption validated that L1210 cells maintain active mitochondrial ATP synthesis, which could be completely inhibited by 1 μM oligomycin, a specific inhibitor of FO-ATP synthase (Supplementary Figs. 7A-C). Unexpectedly, when we treated L1210 cells in the G2 cell cycle phase with 1 μM oligomycin and monitored their growth using the SMR, the cells still proceeded through mitosis and divided symmetrically (Fig. 3A), although the magnitude of the TMRE signal increase in mitosis was reduced (Fig. 3B). To further quantitatively analyze the role of mitochondrial ATP synthesis in mitotic entry and progression, we synchronized cells to G2 using RO-3306, treated the cells with 1 μM oligomycin for 15 min, released the cells to enter mitosis in the presence of oligomycin, and collected samples for cell cycle analysis at different timepoints. Surprisingly, mitochondrial ATP synthesis inhibition had little effect on mitotic entry and the subsequent appearance of G1 cells (Figs. 3C, 3D). Similar results were observed in BaF3 and DT40 lymphocytes (Supplementary Fig. 8). To further examine the extent to which ATP synthesis inhibition influences L1210 cell behaviour, we monitored single-cell mass accumulation (growth) rates using a serial SMR ^15,24^. We observed that oligomycin treatment caused a major reduction in cell growth rates that persisted for several hours (Fig. 3E). Thus, mitochondrial ATP synthesis is required to support cell growth, but not cell division. This finding is consistent with prior observations that mitochondrially localized dominant negative form of CDK1 did not affect G2/M progression despite reducing mitochondrial respiration ^4^, and that cells devoid of mitochondrial DNA and mitochondrial ATP production can proliferate despite significantly reduced growth rates ^25^.

**Fig. 2:**
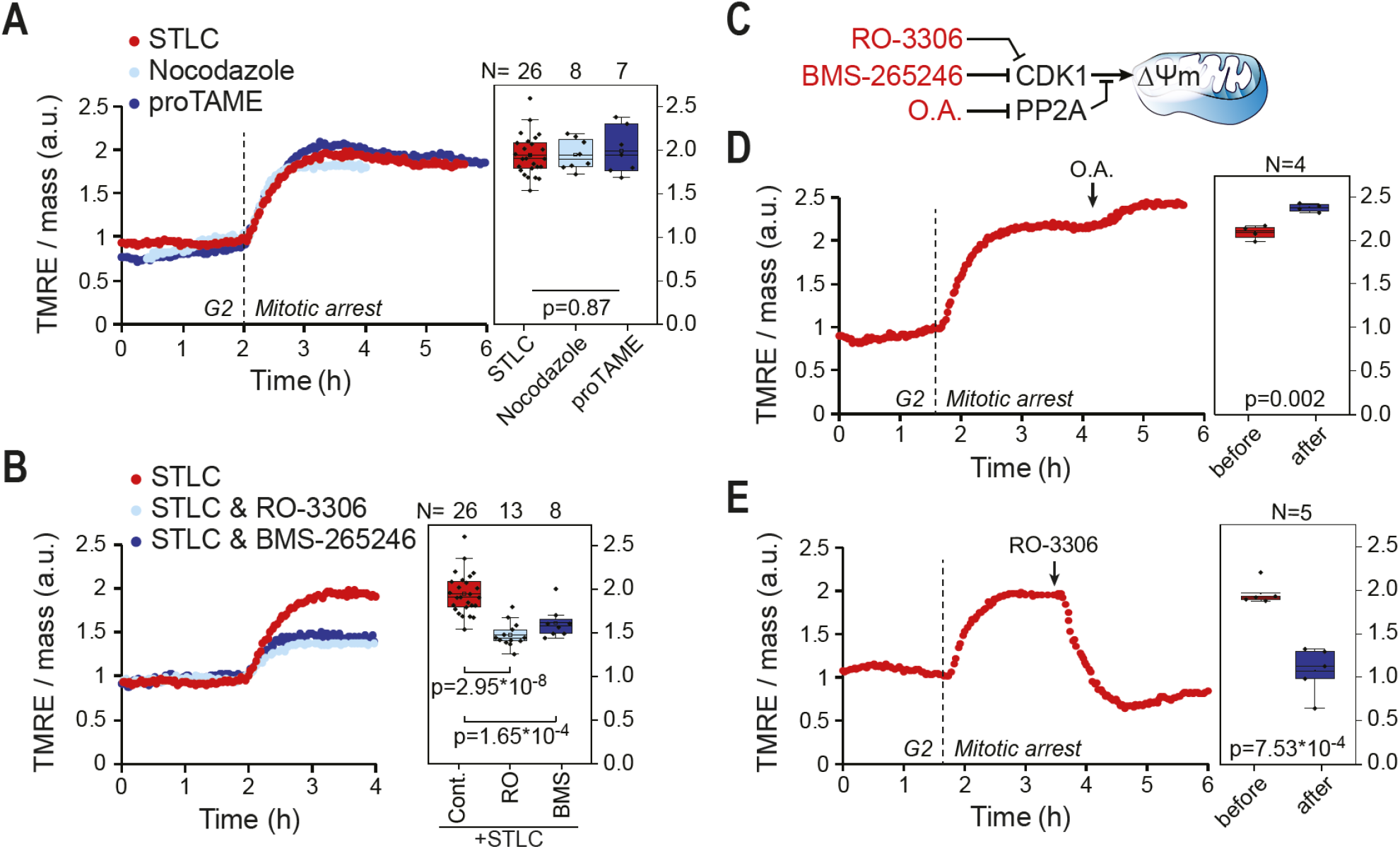
CDK1 drives a switch-like mitochondrial hyperpolarization. **(A)** Mass-normalized TMRE traces for L1210 cells treated with 5 μM STLC (dark blue), 1 μg/ml Nocodazole (light blue) or 15 μM proTAME (red) to induce mitotic arrest. TMRE signal reaches a stable but highly elevated level when mitosis is prolonged, indicative of a metabolic switch in mitosis. Boxplots on the right indicate the level to which TMRE increases during mitotic arrest. p-value was obtained using ANOVA. **(B)** Mass-normalized TMRE traces for L1210 cells treated with 5 μM STLC alone (dark blue), in combination with 1 μM RO-3306 (light blue) or in combination with 400 nM BMS-265246 (red) to partly inhibit CDK1. Partial inhibition of CDK1 reduces the extent of the metabolic switch in mitosis. Boxplots on the right indicate the level to which TMRE increases during mitotic arrest. p-values were obtained using ANOVA followed by Sidakholm test. **(C)** Schematic indicating how the chemical inhibitors (red) affect CDK1 activity. **(D)** Mass-normalized TMRE trace for a L1210 cell treated with 5 μM STLC. Once the cell was arrested in mitosis, 100 nM O.A. was added (black arrow) to the culture media. Boxplots on the right indicate the TMRE level before and after O.A. addition. p-value was obtained using two-tailed Student’s t-test. **(E)** Mass-normalized TMRE trace for a L1210 cell treated with 5 μM STLC. Once the cell was arrested in mitosis, 5 μM RO-3306 was added (black arrow) to the culture media to inhibit CDK1. Boxplots on the right indicate the TMRE level before and after RO-3306 addition.

**Fig. 3:**
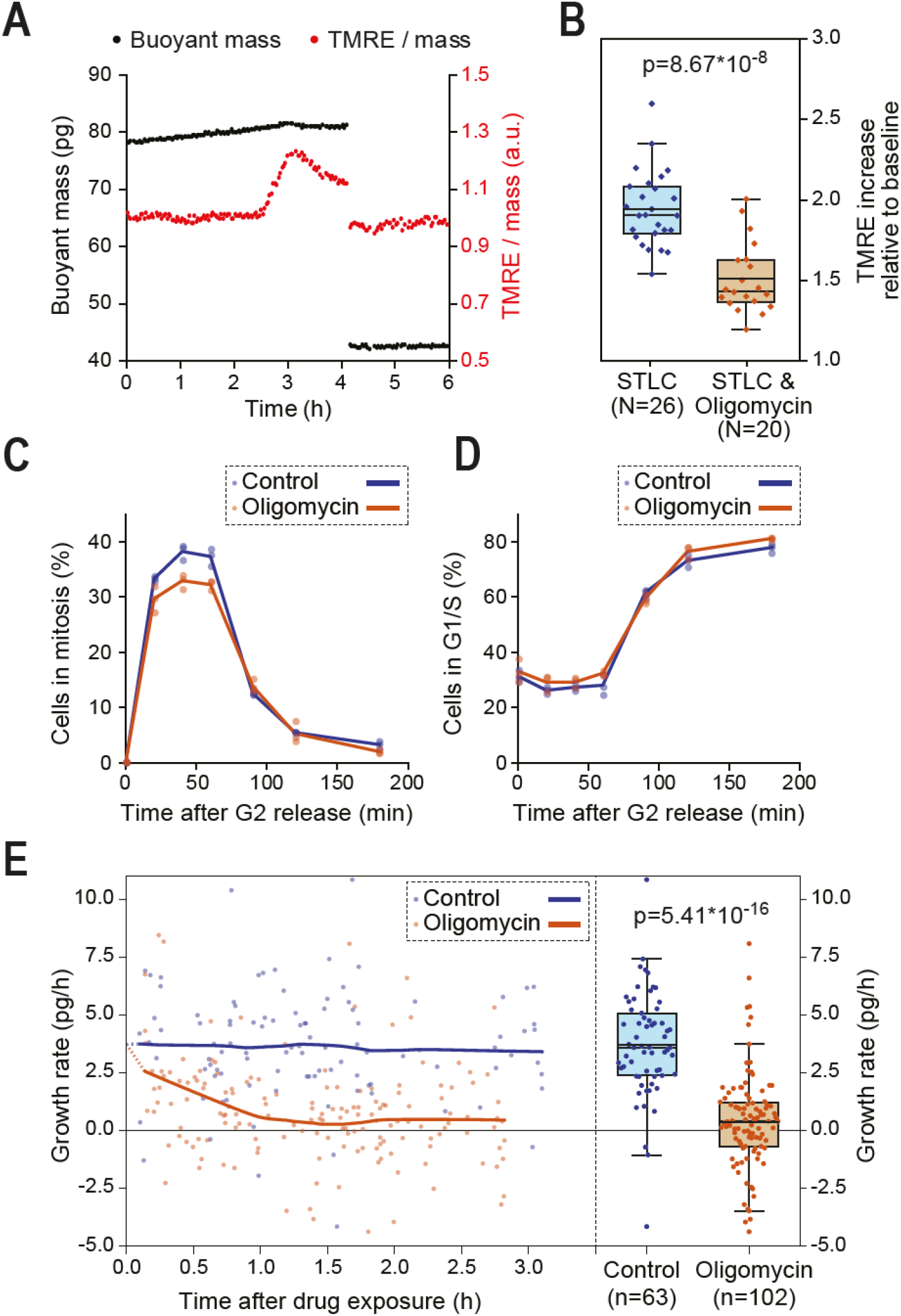
Mitochondrial ATP synthesis is required for cell growth, but not for cell division. **(A)** Buoyant mass (black) and mass-normalized TMRE (red) trace for a L1210 cell around cell division in the presence of 1 μM oligomycin. Despite reduced growth rate, the cell proceeds through mitosis. **(B)** Quantifications of the TMRE increase in mitosis relative to G2 levels in control and 1 μM oligomycin-treated L1210 cells. p-value was obtained using two-tailed Student’s t-test. **(C)** Quantifications of mitotic entry in control and 1 μM oligomycin-treated L1210 cells. Cells were synchronized to G2, released and collected for cell cycle analysis at indicated timepoints. 1 μM oligomycin treatment was started 15 min before release from G2 arrest. Each dot represents a separate culture (n=3). **(D)** Quantifications of mitotic exit (appearance of G1 cells) for samples shown in (C). **(E)** Quantification of mass accumulation (growth) rate in individual control and 1 μM oligomycin-treated cells. Oligomycin blocks cell growth within 1h. Each dot represents the growth rate of a single-cell. Quantifications of the growth rates after 1h drug exposure are shown on the right. p-value was obtained using two-tailed Student’s t-test.

Next, to more directly examine bioenergetics and oxidative stress in mitosis, various fluorescence based metabolic reporters were expressed in both L1210 and BaF3 cells. However, the expression of these exogenous proteins resulted in the loss, or even the reversal of the normal mitotic mitochondrial hyperpolarization (Supplementary Fig. 9, Supplementary note 3), indicating that these genetic tools can bias quantitative analyzes of mitotic mitochondrial bioenergetics in our model system.

As an alternative approach to understand mitotic bioenergetics, we developed a model to derive ATP synthesis rates from the TMRE signal dynamics (Supplementary Fig. 10, Supplementary note 4). First, we converted the TMRE signal to approximate ΔΨm using existing measurements of tetramethylrhodamidine ester dye accumulation and membrane potentials ^26^. Second, we assumed that the voltage across the inner mitochondrial membrane (ΔΨm) is determined by the currents through ATP synthesis (I_ATP_), voltage-dependent leakage ^27^ (I_Leak_) and the ETC (I_ETC_). Third, we modelled the inner mitochondrial membrane as an electrical circuit with voltage (ΔΨm), capacitance (C) and resistances (R) for each one of the currents (Fig. 4A, Supplementary note 4), constituting a simple model that is consistent with the biochemical view of mitochondria ^16,28^. We assumed that the circuit behaves in a switch-like manner between two distinct states whether CDK1 is activated (*CDK1 on*) or inactivated (*CDK1 off*). By fitting our model’s analytical solution to the ΔΨm data we derive RC values, which reflect the time constant of the ΔΨm change (Figs. 4B, 4C, Supplementary notes 5 and 6). We derived these RC values separately for each control and oligomycin-treated cell during *CDK1 on* and *CDK1 off* states (Fig. 4D, Supplementary Fig. 11). Comparing the RC values between control and oligomycin-treated cells, we extracted the resistance of ATP synthase (R_ATP_) during the *CDK1 on* and *CDK1 off* states (Fig. 4B). We found that R_ATP_ is higher during the *CDK1 on* state than during *CDK1 off* state (Supplementary Fig. 11C, Supplementary note 4). In addition, comparing RC values during the *CDK1 on* and *off* states in each control cell revealed that in order for R_ATP_ to increase, R_Leak_ and R_ETC_ are required to cumulatively decrease during the *CDK1 on* state (Supplementary note 7). We then derived the current through ATP synthase (I_ATP_), i.e. the ATP synthesis rate, throughout mitosis using Ohm’s law (I_ATP_=V/R_ATP_) (Supplementary notes 7-9). Surprisingly, our modelling revealed that the mitochondrial ATP synthesis rate is inhibited by 54 % ± 11 % (mean±s.e.m.) during prometaphase and metaphase, when compared to G2 ATP synthesis rates (Figs. 4E, 4F). During anaphase, only a minor increase (<10%) in comparison to G2 levels was observed (Fig. 4E). Overall, this temporal control of ATP synthesis results in 40 % decrease in total mitochondrial ATP production during early mitosis (between G2/M transition and metaphase-anaphase transition) when compared to a situation where mitochondrial ATP synthesis would remain at G2 levels (Fig. 4G).

**Fig. 4:**
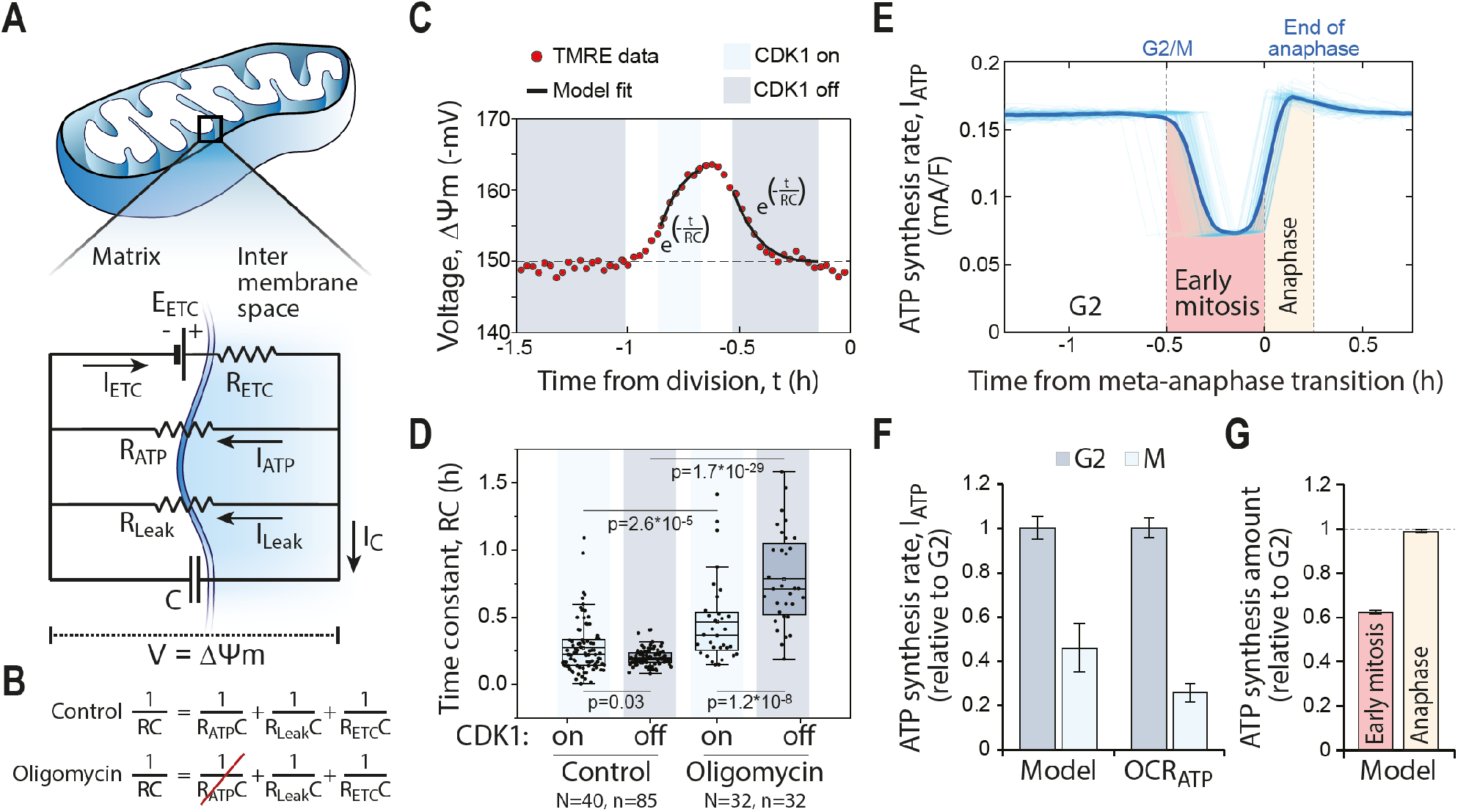
Mitochondrial ATP synthesis is inhibited in mitosis. **(A, B)** Electrical circuit model of mitochondria, where ΔΨm is the voltage across the circuit (*panel A*). The key components controlling ΔΨm (ETC, ATP synthase and leakage), all have their respective resistances (R_ETC_, R_ATP_ and R_Leak_) which together form the total resistance (R) in the circuit (*panel B*). In the presence of oligomycin, the R_ATP_ value increases to infinity. The difference in R values between control and oligomycin-treated cells reflects the R_ATP_ value. **(C)** Model fit to a typical L1210 single-cell TMRE data. TMRE signals were converted to approximate voltages and the period where CDK1 is on (light blue) and off (dark blue) were fitted separately to derive the time constant (RC) values for each period. **(D)** Derived time constant (RC) values for CDK1 on (light blue) and off (dark blue) periods in control and 1 μM oligomycin-treated L1210 cells. p-values were obtained using ANOVA followed by Sidakholm test. **(E)** ATP synthesis rate (I_ATP_) for 85 separate L1210 cells around cell division. Traces of individual cells are drawn with thin opaque lines, whereas the population average is drawn as thick solid line. The colored areas reflect total amounts of ATP synthesized during early mitosis (red) and anaphase (light yellow). **(F)** Quantifications of relative ATP synthesis rates (I_ATP_) in G2 (dark blue) and mitosis (light blue) as modelled on a single-cell level in non-arrested cells (left), and as estimated based on oxygen consumption rates (OCR_ATP_) in cell populations arrested to G2 and mitosis (right). Note that both approaches rely on comparing control and oligomycin treatment conditions to derive ATP synthesis rates, but the model accounts for changes in ΔΨm and does not require cell cycle synchronizations. Data depicts mean ± s.e.m. (n=85-32 for model, n=7-8 for OCR). **(G)** Quantifications of the total amount of mitochondrial ATP synthesis during early mitosis (from G2/M to M/A transition, red) and during anaphase (from M/A transition to end of anaphase where cell elongation is complete, light yellow), when compared to a null-hypothesis where ATP synthesis rate remains at G2 levels throughout mitosis (dashed horizontal line). Data depicts mean ± s.e.m. of the R_ATP_ values (N=40-32, n=85-32).

It is important to recognize the limitations of our approach. Especially, i) the TMRE signal is a proxy for ΔΨm, subject to systematic errors, ii) we assumed that oligomycin only perturbs mitochondrial ATP synthase, and iii) our electric circuit model, while including all major elements of mitochondrial bioenergetics, may oversimplify the dynamics of ATP synthesis. To support our conclusions, we independently quantified the average ATP synthesis rate of prometaphase and G2 arrested cell populations by measuring oligomycin sensitive oxygen consumption (Supplementary Figs. 7D, 7E). Consistently with our modeling approach, this indicated that mitochondrial ATP synthesis rate is significantly reduced in mitotic cells when compared to G2 cells (Fig. 4F). Furthermore, oxygen consumption measurements revealed that mitotically arrested cells have higher leakage than G2 cells (Supplementary Figs. 7D, 7E), consistent with our modeling (Supplementary note 7). In addition, our results on ATP synthesis have little sensitivity to the model-specific parameters, including those used for TMRE-to-ΔΨm conversion (Supplementary Figs. 12-14, Supplementary note 10). Since our modeling relies on comparisons between control and oligomycin-treated cells, any systematic bias that affects TMRE in both samples will not affect our ATP synthesis results.

Our findings are compatible with existing literature. Our observation that ΔΨm can increase in mitosis in the presence of oligomycin (Fig. 3B) is supported by the reported mitotic activation of the ETC ^4,8,9^. Both ETC activation and ATP synthesis inhibition cause ΔΨm to radically increase, and high ΔΨm is known to promote mitochondrial protein import ^9^, reactive oxygen species (ROS) generation ^29^, proton leakage ^27^ and heat production ^30^. Consistently, mitochondrial protein import, ROS levels and cellular heat output have been reported to increase during mitosis in a CDK1-dependent manner ^9,31–33^. Furthermore, the ATP synthesis dynamics we discover can explain the reported ATP level dynamics in mitosis ^3,7,10^.

Overall, our work reveals the previously unknown dynamics of ΔΨm and mitochondrial ATP synthesis during mitosis. Considering that mitochondria are responsible for the majority of cellular ATP synthesis ^11–13^ and that ATP consumption in interphase consumes cellular ATP pools within minutes ^32^, our discovery that mitochondrial ATP synthesis is inhibited during mitosis suggests a much lower rate of ATP consumption during mitosis than previously assumed. Notably, cells maintain ATP at concentrations near 4 millimolar ^7,34^, but most enzymes have Michaelis constants (K_m_) for ATP in the micromolar range. Thus, even a major decrease in cellular ATP levels will not affect enzymatic reaction rates, and the intracellular ATP pools may be able to fulfill the energetic needs of mitosis even in the absence of additional ATP synthesis, as suggested by early work on antephase ^35,36^. The decreased ATP levels may even promote cell division by facilitating cellular reorganization and chromatin condensation ^7,37^.

## Acknowledgements

We would like to thank Zhaoqi Li, Douaa Mugahid and Matthew Vander Heiden for their helpful comments. This research was supported by a Samsung Scholarship (J.H.K.), the Koch Institute Frontier Research Program through the Kathy and Curt Marble Cancer Research Fund (S.R.M.), the MIT Center for Precision Cancer Medicine and National Institute of Health grants P30-CA14051, R01-GM104047 (M.B.Y.), R35-ES028374 (M.B.Y.), and by the Wellcome Trust grant 110275/Z/15/Z (T.P.M.).

## Author contributions

J.H.K., M.A.S. and T.P.M. planned and carried out the experiments. G.K. carried out the modeling. D.L. and T.P.M. generated the reporter cell lines. J.H.K., G.K. and T.P.M. wrote the manuscript with input from all authors. M.B.Y., S.R.M. and T.P.M. supervised the work. T.P.M. conceived the study.

## Supplementary materials

Supplementary notes 1-10

Materials and methods

Supplementary Figs. 1-14

Supplementary references

## References

1 Salazar-Roa, M. & Malumbres, M. Fueling the Cell Division Cycle. Trends Cell Biol 27, 69–81, doi:10.1016/j.tcb.2016.08.009 (2017).

2 Lopez-Mejia, I. C. & Fajas, L. Cell cycle regulation of mitochondrial function. Curr Opin Cell Biol 33, 19–25, doi:10.1016/j.ceb.2014.10.006 (2015).

3 Zhao, H. et al. AMPK-mediated activation of MCU stimulates mitochondrial Ca(2+) entry to promote mitotic progression. Nat Cell Biol 21, 476–486, doi:10.1038/s41556-019-0296-3 (2019).

4 Wang, Z. et al. Cyclin B1/Cdk1 coordinates mitochondrial respiration for cell-cycle G2/M progression. Dev Cell 29, 217–232, doi:10.1016/j.devcel.2014.03.012 (2014).

5 Li, Z. Y., Ji, X. M., Wang, D. M., Liu, J. J. & Zhang, X. Autophagic flux is highly active in early mitosis and differentially regulated throughout the cell cycle. Oncotarget 7, 39705–39718, doi:10.18632/oncotarget.9451 (2016).

6 Liu, L., Xie, R., Nguyen, S., Ye, M. & McKeehan, W. L. Robust autophagy/mitophagy persists during mitosis. Cell Cycle 8, 1616–1620, doi:10.4161/cc.8.10.8577 (2009).

7 Maeshima, K. et al. A Transient Rise in Free Mg(2+) Ions Released from ATP-Mg Hydrolysis Contributes to Mitotic Chromosome Condensation. Curr Biol 28, 444–451 e446, doi:10.1016/j.cub.2017.12.035 (2018).

8 Domenech, E. et al. AMPK and PFKFB3 mediate glycolysis and survival in response to mitophagy during mitotic arrest. Nat Cell Biol 17, 1304–1316, doi:10.1038/ncb3231 (2015).

9 Harbauer, A. B. et al. Mitochondria. Cell cycle-dependent regulation of mitochondrial preprotein translocase. Science 346, 1109–1113, doi:10.1126/science.1261253 (2014).

10 Chin, B. & Bernstein, I. A. Adenosine triphosphate and synchronous mitosis in Physarum polycephalum. J Bacteriol 96, 330–337 (1968).

11 Moreno-Sanchez, R. et al. Who controls the ATP supply in cancer cells? Biochemistry lessons to understand cancer energy metabolism. Int J Biochem Cell Biol 50, 10–23, doi:10.1016/j.biocel.2014.01.025 (2014).

12 Zu, X. L. & Guppy, M. Cancer metabolism: facts, fantasy, and fiction. Biochem Biophys Res Commun 313, 459–465 (2004).

13 Moreno-Sanchez, R., Rodriguez-Enriquez, S., Marin-Hernandez, A. & Saavedra, E. Energy metabolism in tumor cells. FEBS J 274, 1393–1418, doi:10.1111/j.1742-4658.2007.05686.x (2007).

14 Kang, J. H. et al. Noninvasive monitoring of single-cell mechanics by acoustic scattering. Nat Methods 16, 263–269, doi:10.1038/s41592-019-0326-x (2019).

15 Miettinen, T. P., Kang, J. H., Yang, L. F. & Manalis, S. R. Mammalian cell growth dynamics in mitosis. Elife 8, doi:10.7554/eLife.44700 (2019).

16 Nicholls, D. G. & Ward, M. W. Mitochondrial membrane potential and neuronal glutamate excitotoxicity: mortality and millivolts. Trends Neurosci 23, 166–174 (2000).

17 Son, S. et al. Resonant microchannel volume and mass measurements show that suspended cells swell during mitosis. J Cell Biol 211, 757–763, doi:10.1083/jcb.201505058 (2015).

18 Zlotek-Zlotkiewicz, E., Monnier, S., Cappello, G., Le Berre, M. & Piel, M. Optical volume and mass measurements show that mammalian cells swell during mitosis. J Cell Biol 211, 765–774, doi:10.1083/jcb.201505056 (2015).

19 Son, S. et al. Direct observation of mammalian cell growth and size regulation. Nat Methods 9, 910–912, doi:10.1038/nmeth.2133 (2012).

20 Marteil, G. et al. Proteomics reveals a switch in CDK1-associated proteins upon M-phase exit during the Xenopus laevis oocyte to embryo transition. Int J Biochem Cell Biol 44, 53–64, doi:10.1016/j.biocel.2011.09.003 (2012).

21 Lindqvist, A., Rodriguez-Bravo, V. & Medema, R. H. The decision to enter mitosis: feedback and redundancy in the mitotic entry network. J Cell Biol 185, 193–202, doi:10.1083/jcb.200812045 (2009).

22 Gavet, O. & Pines, J. Progressive activation of CyclinB1-Cdk1 coordinates entry to mitosis. Dev Cell 18, 533–543, doi:10.1016/j.devcel.2010.02.013 (2010).

23 Hegarat, N., Rata, S. & Hochegger, H. Bistability of mitotic entry and exit switches during open mitosis in mammalian cells. Bioessays 38, 627–643, doi:10.1002/bies.201600057 (2016).

24 Cermak, N. et al. High-throughput measurement of single-cell growth rates using serial microfluidic mass sensor arrays. Nat Biotechnol 34, 1052–1059, doi:10.1038/nbt.3666 (2016).

25 Miettinen, T. P. et al. Identification of transcriptional and metabolic programs related to mammalian cell size. Curr Biol 24, 598–608, doi:10.1016/j.cub.2014.01.071 (2014).

26 Gerencser, A. A. et al. Quantitative measurement of mitochondrial membrane potential in cultured cells: calcium-induced de- and hyperpolarization of neuronal mitochondria. J Physiol 590, 2845–2871, doi:10.1113/jphysiol.2012.228387 (2012).

27 Nobes, C. D., Brown, G. C., Olive, P. N. & Brand, M. D. Non-ohmic proton conductance of the mitochondrial inner membrane in hepatocytes. J Biol Chem 265, 12903–12909 (1990).

28 Nicholls, D. G. Mitochondrial membrane potential and aging. Aging Cell 3, 35–40 (2004).

29 Murphy, M. P. How mitochondria produce reactive oxygen species. Biochem J 417, 1–13, doi:10.1042/BJ20081386 (2009).

30 Picard, M., McEwen, B. S., Epel, E. S. & Sandi, C. An energetic view of stress: Focus on mitochondria. Front Neuroendocrinol 49, 72–85, doi:10.1016/j.yfrne.2018.01.001 (2018).

31 Patterson, J. C. et al. ROS and Oxidative Stress Are Elevated in Mitosis during Asynchronous Cell Cycle Progression and Are Exacerbated by Mitotic Arrest. Cell Syst 8, 163–167 e162, doi:10.1016/j.cels.2019.01.005 (2019).

32 Rodenfels, J., Neugebauer, K. M. & Howard, J. Heat Oscillations Driven by the Embryonic Cell Cycle Reveal the Energetic Costs of Signaling. Dev Cell 48, 646–658 e646, doi:10.1016/j.devcel.2018.12.024 (2019).

33 Lim, J. M., Lee, K. S., Woo, H. A., Kang, D. & Rhee, S. G. Control of the pericentrosomal H2O2 level by peroxiredoxin I is critical for mitotic progression. J Cell Biol 210, 23–33, doi:10.1083/jcb.201412068 (2015).

34 Traut, T. W. Physiological concentrations of purines and pyrimidines. Mol Cell Biochem 140, 1–22 (1994).

35 Gelfant, S. The energy requirements for mitosis. Ann N Y Acad Sci 90, 536–549 (1960).

36 Swann, M. M. The Control of Cell Division - a Review .1. General Mechanisms. Cancer Res 17, 727–757 (1957).

37 Patel, A. et al. ATP as a biological hydrotrope. Science 356, 753–756, doi:10.1126/science.aaf6846 (2017).

