## Supplementary materials for "Time-dynamics of mitochondrial membrane potential reveal an inhibition of ATP synthesis in mitosis"

**This PDF file includes:**

Table of Contents
Supplementary Notes 1-10
Materials and Methods
Supplementary Figures 1-14
Supplementary References

#### Supplementary Note 1. Non-invasive, continuous, high-resolution monitoring of mass normalized fluorescence signals in single-cells

We wanted to study the cell cycle dependent behavior of mitochondrial bioenergetics. To do this, we sought for a method that i) allows continuous monitoring of single-cell fluorescence reporters, such as mitochondrial membrane potential (ΔΨm) sensitive fluorescent probes, ii) enables fluorescence measurements with high temporal resolution and precision, iii) is non-invasive by minimizing phototoxicity, iv) allows cell size normalization of the fluorescence signals. To achieve this, we utilized a microfluidic mass sensor called the suspended microchannel resonator (SMR). SMR quantifies the buoyant mass of a single-cell non-invasively as the cell flows through a vibrating cantilever [1-4]. Using a hydrodynamic trapping approach, the cell can be repeatedly measured over multiple cell cycles, where one of the daughter cells is randomly discarded and the other is monitored following each division. We coupled the SMR to a fluorescence detection setup, where the excitation laser would only be turned on for a brief period during which the cell passes through the laser and the total fluorescence emission is quantified using photomultiplier tubes (Supplementary Figs. 1A, 1B, Materials and Methods). This approach allowed us to monitor mass normalized fluorescence signals of single-cells with a typical temporal resolution of 2 min over multiple cell cycles without perturbing normal cell growth (Fig. 1A) [2,3,5]. Detailed characterization of our mass measurement precision (measurement error < 0.1 pg for a typical L1210 cell) is previously published [3] and details of our fluorescence measurement precision can be found below (Supplementary note 2).

#### Supplementary Note 2. TMRE as a probe for ΔΨm

We selected TMRE as our main proxy for ΔΨm due to TMRE's low mitochondrial binding, low electron transport chain inhibition and fast equilibrium [6-10]. As lipophilic cation, TMRE accumulates in mitochondria based on the negative charge in mitochondrial matrix. However, the cytosol also has a slightly negative charge compared to the extracellular media. Thus, while TMRE accumulates primarily to mitochondria and the total TMRE signal reflects primarily ΔΨm, some TMRE can also accumulate in the cytosol and bias the total TMRE signal intensity as a probe for ΔΨm [7-9,11,12]. Below, we show in L1210 cells that i) mitotic increase in TMRE signal reflects an increased ΔΨm, ii) mitotic changes in plasma membrane potential (ΔΨp) or cell volume do not significantly contribute to the TMRE signal, and iii) TMRE diffusion speed or our fluorescence detection accuracy do not bias our quantifications of TMRE dynamics.

***Non-quenching and quenching mode experiments with ΔΨm responsive probes***

In all our experiments with TMRE, we used TMRE in non-quenching concentrations (10 nM). In a non-quenching mode, increased ΔΨm results in increased TMRE accumulation in mitochondria and, consequently, increased TMRE fluorescence from the cells. We validated this TMRE behavior by increasing ΔΨm using oligomycin (an inhibitor of ATP synthase) and by reducing ΔΨm using FCCP (a protonophore) (Supplementary Fig. 2A).

ΔΨm responsive probes, such as TMRE or Rhod123, can also be used in very high concentrations where the probes switch to quenching mode [6]. In the quenching mode, the ΔΨm probe accumulates to mitochondria in such a high concentration that the fluorescence signal is quenched. Thus, increased ΔΨm results in increased probe accumulation in mitochondria which further quenches the probe fluorescence, resulting in decreased total fluorescence from the cells. We validated this behavior for 10 μM Rho123 using oligomycin and FCCP (Supplementary Fig. 2B). Notably, in the quenching mode experiments, the cell culture media does not have Rhod123. Thus, any changes in the total cell fluorescence, as seen in mitosis (Fig. 1B), cannot be explained by artificial uptake of the probe or by changes in plasma membrane potential. Together with the TMRE measurements in non-quenching mode (Fig. 1A), these results indicate increased ΔΨm in mitosis.

***Influence of cell mass, volume and mitochondrial content on TMRE signal in mitosis***

Changes in cell size and mitochondrial content can affect the TMRE accumulation to the cell and, consequently, TMRE signal. We normalized our TMRE measurements to the buoyant mass of a cell, enabling us to account for changes in cell mass observed throughout the cell cycle (Fig. 1A). Importantly, in early mitosis, when TMRE signal increases, cells undergo mitotic cell swelling, which increases cell volume, but has little influence on buoyant mass [1,3,13]. To exclude the possibility that this volume increase is responsible for the TMRE increase, we inhibited the mitotic cell swelling, and consequently the volume increase, using EIPA [1]. By doing so, we observed that the mitotic TMRE increase was not dependent on the volume increase (Supplementary Fig. 4). This may reflect the relatively small cytosolic volume present in L1210 lymphocytes, which reduces the influence that cytosolic volume has on TMRE accumulation. Furthermore, FL5.12 cells, which also increase their volume in mitosis [1], did not display TMRE increase in mitosis (Supplementary Fig. 3A). We also examined how mitochondrial content changes during mitosis by monitoring of the ΔΨm-insensitive MitoTracker Green probe. As mitochondrial content is known to scale with cell size [14-16], we normalized the MitoTracker Green signal to cell's buoyant mass. The mass-normalized MitoTracker Green signal did not change during the mitotic mitochondrial hyperpolarization (Fig. 1C), indicating that mitochondrial volume does not radically alter during mitosis. Thus, changes in cytosolic or mitochondrial volumes don't explain the TMRE behavior in mitosis.

***Influence of ΔΨp on TMRE signal in mitosis***

Non-quenching mode experiments with ΔΨm responsive probes can be affected by changes in ΔΨp. To assess how ΔΨp changes in mitosis, we monitored the ΔΨp responsive probe DiBAC$_4$(3) and normalized its fluorescence signal to the cell's buoyant mass. We observed a decrease in DiBAC$_4$(3) signal during early mitosis (Supplementary Fig. 4), suggesting an increase in ΔΨp. However, the ΔΨp increase occurred earlier than TMRE increase (Supplementary Fig. 4A), similarly to the mitotic cell swelling (Fig. 1E), suggesting that the ΔΨp change has only limited influence on TMRE signal. We then inhibited mitotic cell swelling using EIPA and observed that the EIPA treatment reduced the DiBAC$_4$(3) change (Supplementary Figs. 4A, 4B). Importantly, inhibition of mitotic swelling, and thus of ΔΨp increase, did not reduce the TMRE increase in mitosis (Supplementary Fig. 4C). Thus, the extent to which plasma membrane hyperpolarizes is not enough to significantly affect TMRE signal. This is consistent with our other observations that i) quenching mode experiments with Rhod123 display increased ΔΨm (Fig. 1B), and ii) the inhibition of mitochondrial ATP synthase with oligomycin significantly influences the mitotic TMRE dynamics (Fig. 4D). In conclusion, while changes in ΔΨp will always influence TMRE signal, the mitotic TMRE increase in L1210 cells reflects primarily changes in ΔΨm.

***Characterizing TMRE diffusion dynamics and TMRE detection accuracy***

We also considered the limits of our approach in detecting the precise kinetics of mitotic TMRE increase and decrease. Our observations of TMRE dynamics are limited by our measurement precision and frequency, and the diffusion rate of TMRE in response to changes in ΔΨm. We first quantified our TMRE measurement precision. To account for both the technical error in our measurements and the biological variability in mitochondrial localization, which can also influence the total fluorescence detected, we labelled mitochondria using MitoTracker Red CMXRos, fixed the cells, and repeatedly measured individual cells to quantify the error for each fixed cell. This indicated that our measurement error is dependent on the total fluorescence intensity (Supplementary Fig. 1C) and for a typical mitotic L1210 cell our TMRE measurement error is approximately 2% (Supplementary Fig. 1D). The typical frequency of our TMRE measurements when monitoring live cells in mitosis was 2 min. Thus, our temporal resolution was approximately 4 min according to the Nyquist rate. In conclusion, TMRE signal change during the mitotic mitochondrial hyperpolarization (~50% increase in TMRE signal over ~30 min period) is an order of magnitude larger that our TMRE measurement precision and temporal resolution.

Next, we assessed if TMRE diffusion speed is limiting our ability to detect the full dynamics of mitotic mitochondrial hyperpolarization. TMRE is known to diffuse across cell membranes rapidly, reaching equilibrium in minute timescales, especially in small cells with high surface-to-volume ratio [7,8,17,18]. We took two separate experimental approaches to validate that TMRE diffusion rate is fast enough to capture the ΔΨm dynamics during mitosis. First, we monitored mitotic TMRE behavior in the presence of 1 μM tetraphenylborate (TPB). TPB is known to facilitate the diffusion of tetramethylrhodamidine esters through the plasma membrane, thus increasing the rate at which equilibrium is reached [7,8]. We did not observe changes in mitotic TMRE dynamics in the presence of TPB (Supplementary Fig. 2C). Second, we monitored TMRE signal decay rate on a population level following mitochondrial uncoupling using 10 μM FCCP

(Supplementary Fig. 2D). We used this data to quantify exponential time constants for TMRE signal loss (i.e. TMRE diffusion speed out of the cells). We then compared the time constants obtained after FCCP treatment to time constants obtained by fitting an exponential decay to the TMRE depolarization observed in late mitosis (Fig. 4D). Depending on the data fitting used to derive the time constant values, TMRE diffusion speed was 1 or 2 orders of magnitude faster than the TMRE signal change in mitosis (Supplementary Fig. 2E). Thus, in the mitosis of L1210 cells, TMRE signal is a reliable proxy for capturing the ΔΨm dynamics.

#### Supplementary Note 3. Expression of exogenous proteins alters mitochondrial metabolism in mitosis

We aimed to characterize metabolic changes during mitosis by monitoring genetic FRET based metabolic sensors throughout mitosis. We generated L1210 and BaF3 cells that stably express an ATP reporter (A-Team), $Ca^{2+}$ reporter (GCaMP3), reactive oxygen species reporter (roGFP2-Orp1) or glutathione redox potential reporter (Grx1-roGFP2) [19-23] localized to either cytosol or mitochondria. To enable reliable detection of the weak fluorescence signals, all constructs were expressed under a strong (CMV) promoter. Surprisingly, expression of all of these constructs removed or even reversed the TMRE increase observed in mitosis (Supplementary Fig. 9A). Furthermore, even the FUCCI L1210 cells, which express mAG-Geminin under the endogenous promoter, did not display as extensive TMRE increase in mitosis as WT L1210 cells (Supplementary Figs. 9B, 9C). Based on these observations, we concluded that expressing high levels of exogenous proteins alters mitochondrial metabolism in mitosis. Thus, the mitotic mitochondrial bioenergetics of wild-type L1210 cells cannot be quantitatively analyzed using genetic reporters.

#### Supplementary Note 4. Modeling mitochondria as an electrical circuit

Our experimental evidence shows that the mitochondrial hyperpolarization in mitosis takes place during prophase, prometaphase and metaphase (Fig. 1), which correlates with the timing of CDK1 activity [24]. Consistently, we show that CDK1 activity is required for the mitochondrial hyperpolarization (Fig. 2). Importantly, CyclinB-CDK1 activity is regulated in a switch-like manner that causes the system to operate in fully active or inactive state with brief transition periods between these two stages [24-26]. Consistently, our data shows that arresting cells in metaphase, where CDK1 is active, results in a new, stable level of mitochondrial polarization (Fig. 2A). Together, these results indicate that the cells switch their metabolic state when passing through the G2/M transition (CDK1 is on) and revert back to the original state when passing through the metaphase/anaphase transition (CKD1 is off).

We reasoned that this biphasic behavior in mitochondrial membrane potential control would allow us to model and quantify mitochondrial bioenergetics in mitosis. To this end, we generated an electrical circuit model of mitochondria (Fig. 4A) which is consistent with the common 'electrical circuit' view of mitochondria [9,18,27,28]. The electrical circuit model focuses on ΔΨm as the ATP synthesis driving force, excluding the pH gradient component of the total proton motive force (Δp). Note that in mammalian cell mitochondria ΔΨm is the dominant component of Δp and ΔΨm is also considered indispensable for ATP synthesis [9,12,18,28-30].

Our model is formed of multiple components. The voltage ($V$) across the circuit represents ΔΨm. The voltage source ($E_{ETC}$) with internal resistance ($R_{ETC}$), which together form a single 'battery', represent the electron transport chain (ETC). Note that we assumed the $E_{ETC}$ to have a voltage, which is higher than what can be generated across the inner mitochondrial membrane. Thus, this 'battery' that represents the ETC generates a current $I_{ETC}$ that flows only in one direction (Fig. 4A). Thus, the current $I_{ETC}$ representing the ETC activity is regulated solely by the $R_{ETC}$. Note that $I_{ETC}$ also represents the transport of other charged ions or molecules that increase the voltage across the inner mitochondrial membrane. We formulated our model so that charge through this 'battery' can only move to one direction, thus excluding potential proton leakage through the 'battery'. Notably, a similar circuit model could also be generated using a current source instead of the voltage source coupled to a resistance. While simple and elegant, a current source does not reflect the true physical nature of the ETC, as its activity is independent of the voltage across the circuit. In contrast, the ETC's proton pumping activity depends on the ΔΨm due to the increasing electron leak at high membrane potentials [31,32]. The ATP synthase is represented in our model as a resistance $R_{ATP}$ and the leakage across the membrane is represented as $R_{LEAK}$. The currents through these components are $I_{ATP}$ and $I_{LEAK}$, respectively. Note that $I_{LEAK}$ also represents the transport of other charged ions or molecules that decrease the voltage across the inner mitochondrial membrane without producing ATP. Both $R_{ATP}$ and $R_{LEAK}$ can be controlled by the cell. To account for the fact that mitochondria hyperpolarize not immediately but within some time scale from the activation of the switch-like activity of CDK1, our electrical model also includes a capacitance ($C$). As the mitochondrial membrane composition or the area of the heavily folded inner mitochondrial membrane are unlikely to change rapidly, we assumed that the capacitance remains constant throughout mitosis.

The underlying principle of our model is to separate mitosis in to two regions, $\mathrm{CDK1_{off}}$ and $\mathrm{CDK1_{on}}$, and assumed that all the R values and $E_{ETC}$ can change between $\mathrm{CDK1_{off}}$ and $\mathrm{CDK1_{on}}$ states. Then, by comparing the total resistance of the circuit in the presence and absence of oligomycin, we can derive $R_{ATP}$ for each region and circumvent dealing with other parameters in the model, assuming that oligomycin only affects ATP synthase (i.e. oligomycin does not change $R_{ETC}$ or $R_{Leak}$). Notably, this approach can be compared to oxygen consumption-based estimation of mitochondrial ATP synthesis, such as the Seahorse Extracellular Flux Analyzer, which rely on similar comparisons between control and oligomycin treated conditions [27]. Furthermore, our model accounts for the well-known non-ohmic ΔΨm-dependent leakage [28,32,33] when deriving ATP synthesis rate, $I_{ATP}$.

#### Supplementary Note 5. Derivation of analytical solution to electric circuit model

Applying Kirchhoff's first law into our electric circuit (Fig. 4A), i.e. equaling the currents entering and exiting a node, we obtain:

$$I_{ETC} = I_{ATP} + I_{Leak} + I_C \tag{1}$$

Applying Ohm's law and capacitor equation to write the currents in terms of Voltage, we obtain:

$$\frac{E_{ETC} - V}{R_{ETC}} = \frac{V}{R_{ATP}} + \frac{V}{R_{Leak}} + C\frac{dV}{dt} \tag{2}$$

Rearranging equation (2), we obtain:

$$\frac{dV}{dt} + \frac{V}{R_{ATP}C} + \frac{V}{R_{Leak}C} + \frac{V}{R_{ETC}C} = \frac{E_{ETC}}{R_{ETC}C} \tag{3}$$

In equation (3), we set expressions:

$$\frac{1}{R} = \frac{1}{R_{ATP}} + \frac{1}{R_{Leak}} + \frac{1}{R_{ETC}} \tag{4}$$

$$I_S = \frac{E_{ETC}}{R_{ETC}} \tag{5}$$

Inserting expressions (4,5), into equation (3) we obtain:

$$\frac{dV}{dt} + \frac{V}{RC} = \frac{I_S}{C} \tag{6}$$

Equation (6) is a 1st order inhomogeneous ordinary differential equation with homogeneous and partial solutions:

$$V_h(t) = c\, e^{-\frac{t}{RC}} \tag{7}$$

$$V_p = I_S R \tag{8}$$

We obtain the full solution which is the sum of the homogeneous and partial solutions and includes a constant $c$ specified by the initial conditions:

$$V(t) = c\, e^{-\frac{t}{RC}} + I_S R \tag{9}$$

At $t = t_1$ when CDK1 turns on, we have $V(t_1) = V_1$ , $I_S R > V_0$ and $RC = RC_{CDK1\,on}$, thus we obtain:

$$V(t) = (V_1 - I_S R)\, e^{-\frac{t-t_1}{RC_{CDK1\,on}}} + I_S R \tag{10}$$

At $t = t_3$ when CKD1 is just turning off, we have $V(t_3) = V_3$, $I_S R = V_0$ and $RC = RC_{CDK1\,off}$, thus we obtain:

$$V(t) = (V_3 - V_0)\, e^{-\frac{t-t_3}{RC_{CDK1\,off}}} + V_0 \tag{11}$$

Given that CKD1 is on at $t = [t_1, t_2]$ and off at $t = [t_3, t_4]$ we compile equations (10,11) and write:

$$V(t) = \begin{cases} (V_1 - I_S R) e^{-\frac{t-t_1}{RC_{CDK1\,on}}} + I_S R \;, & t_1 \le t \le t_2 \\ (V_3 - V_0)\, e^{-\frac{t-t_3}{RC_{CDK1\,off}}} + V_0 \;, & t_3 \le t \le t_4 \end{cases} \tag{12}$$

#### Supplementary Note 6. Obtaining time constants (RCs) by fitting analytical solutions to experimental data

##### *Converting TMRE to voltage*

We first normalized the experimental data of TMRE/mass signal, so that each cell has the same TMRE/mass baseline (Supplementary Fig. 11A). To identify the start $(t = t_0)$ and end of mitosis $(t = t_4)$ where we fit our model, we applied a smoothing filter on the normalized TMRE/mass signal (Supplementary Fig. 11B). Next, we converted the normalized TMRE/mass signal to approximate voltage (Supplementary Fig. 11C) using existing calibration curves between tetramethylrhodamidine ester fluorescence signals and membrane potentials [11]. The conversion was done using the following formula: $V(mV) = (TMRE/c_1)/c_2$, where $c_1 = 29.109$ and $c_2 = 0.0215$. For control cells we set $V_0 = -150\,mV$, which is a typical ΔΨm for many cell types [8,9,11,18,34]. For oligomycin treated cells we set $V_0 = -167.8\,mV$, which was based on the TMRE-to-voltage scaling and the measured TMRE increase in L1210 cells after a treatment with oligomycin (Supplementary Fig. 2A). Note, that while this TMRE-to-voltage conversion provides only estimates of the true ΔΨm, our final modeling results were not sensitive to the values used for TMRE-to-voltage scaling (see Supplementary note 10).

##### *Defining fitting regions*

We divided the voltage traces into regions where CDK1 is on $(t_1 \leq t \leq t_2)$ and off $(t_3 \leq t \leq t_4)$ (Supplementary Fig. 11D). First, we approximated the duration $\Delta t$ that L1210 cells take to turn off CDK1. As CDK1 activity is lost at metaphase-anaphase transition due to the APC/C driven degradation of Cyclin B [24], we estimated the duration of this transition by quantifying the duration that L1210 cells took to degrade Geminin, another protein degraded at metaphase-anaphase transition due to APC/C activity (Supplementary Fig. 6). We then used this duration $\Delta t = 8.57 min$ to exclude the parts of mitosis where CDK1 activity is not fully on or off from our analysis. Note that the dynamics of CDK1 turning on and off have been shown to be comparable [24]. Thus, we defined the start of the region where CDK1 is fully on $(t = t_1)$ as $(t_1 = t_0 + \Delta t)$. Next, we identified where CDK1 is turned off. The end of the period where CDK1 is on $(t = t_2)$(i.e. metaphase-anaphase transition) is expected to be near the occurrence of maximal voltage $(t = t_f)$ (Fig. 1G), but the precise location of maximal voltage can be affected by noise in

TMRE signal. Thus, we scanned the region around $t_f$ to find the most optimal fitting. For every dataset for a single cell, we took eleven candidate values $t_2 = t_f - Split\ \Delta t$ (Supplementary Fig. 11D) where $Split = 1, 11, 21, \ldots, 91\ or\ 99$ %. For each one of these $Split$ values, we fitted the CDK1 on $(t = [t_1, t_2])$ and CDK1 off periods $(t = [t_3, t_4])$ with analytical solutions from equations (10) and (11) correspondingly, excluding the $\Delta t$ regions where CDK1 activity is changing. To provide the best fit to the data (Fig. 4C, Supplementary Fig. 11D), we applied the Trust Region Reflective Algorithm [35] with optimization residuals of tolerance values $10^{-6}$. For the time interval $t = [t_1, t_2]$ where CDK1 is turned on, we obtained optimum values for two parameters $I_S R, RC_{CDK1\,on}$. For the time interval $t = [t_3, t_4]$ where CDK1 is turned off, we obtained the optimal value for the parameter $RC_{CDK1\,off}$. Thus, each $Split$ value yields its own $RC_{CDK1\,on}$ and $RC_{CDK1\,off}$ values with R-squared values $R^2_{CDK1\,on}$ and $R^2_{CDK1\,off}$ describing the quality of the fit. For the modeling, we utilized the Split value that provides highest combined R-squared for $RC_{CDK1\,on}$ and $RC_{CDK1\,off}$ (Supplementary Fig. 11E). The same methodology is applied to both control and oligomycin treated cells with the only difference concerning the baseline voltage as described before. Note that oligomycin had little effect on the mitotic progression (Figs. 3C, 3D), thus allowing us to use the same $\Delta t$ as for control. Thus, for both control and oligomycin treated cells we obtained the time constant $RC_{CDK1\,on}$ and $RC_{CDK1\,off}$ values (Fig. 4D) by fitting or model solution to the regions where CDK1 is on and off, respectively.

***Data exclusion from the model***

It is important to note that we kept datasets of single cells when they satisfy two criteria. First, our model needs to yield a physiologically relevant parameters, in particular the $I_S R$ value (maximum projected voltage of hyperpolarization). We excluded any control and oligomycin fits that yielded $I_S R$ value above -203mV , as experiments with isolated mitochondria have indicated that mitochondrial membrane potential cannot increase beyond values of approximately -225 mV due to the non-ohmic proton conductance [30,36-38] and live cell experiments have suggested lower ΔΨm limits [27,33]. Second, we assumed that when CDK1 is turned off (Equation 11) the voltage returns to baseline levels similar to those observed before the mitotic mitochondrial hyperpolarization. Thus, the TMRE signal (voltage) needs to establish a clear new baseline before the abscission of daughter cells. Therefore, we excluded cells that divided before a new baseline was established. In total these exclusion criteria resulted in using 85 cells of the measured 124 control cells and 32 out of the 38 measured oligomycin cells.

#### Supplementary Note 7. Calculating ATP current ($I_{ATP}$) from time constants (RCs)

##### *Approach for calculating $I_{ATP}$ currents*

To calculate the current $I_{ATP}$ we used Ohm's law using the voltage and the $R_{ATP}C$ values for each case of $CDK1\ on$ and $CDK1\ off$:

$$\frac{I_{ATP,CDK1\,off}}{C} = V\left(\frac{1}{R_{ATP}C}\right)_{CDK1\,off} \tag{13}$$

$$\frac{I_{ATP,CDK1\,on}}{C} = V\left(\frac{1}{R_{ATP}C}\right)_{CDK1\,on} \tag{14}$$

Note that we calculated the currents per capacitance $C$ which we assumed to be constant. We used the voltage $V$ values converted from the experimentally measured TMRE. To calculate $R_{ATP}C$ for each case of $CDK1\ on$ and $CDK1\ off$ we used the mean values of the time constants $RC_{CDK1\,on}$ and $RC_{CDK1\,off}$. Note that $RC_{CDK1\,on}$ and $RC_{CDK1\,off}$ originate from single-cell experiments, but the derivation of $R_{ATP}C$ requires comparisons between control and oligomycin conditions. Therefore, we calculated two mean values $R_{ATP}C$, one for $CDK1\ on$ and one for $CDK1\ off$. We then applied these values to all control cell voltage traces in order to calculate $I_{ATP}$ based on equations (13,14), while taking in to account the non-ohmic scaling of leakage with increasing voltage. Below, for simplicity in writing for the rest of this section, we omitted the mention of subscripts $CDK1\ on$ and $CDK1\ off$. Moreover, we referred to mean value as $RC$ with superscripts of $C$ for control, $O$ for oligomycin treated. In addition, we used subscripts of $ATP, Leak, ETC$ for different components of the electrical circuit, consistently with equations (2-5) and main text (Fig. 4A).

##### *$I_{ATP}$ for CDK1$_{off}$*

For control cells, we apply equation (4):

$$\left(\frac{1}{RC}\right)^C = \left(\frac{1}{R_{ATP}C}\right)^C + \left(\frac{1}{R_{Leak}C}\right)^C + \left(\frac{1}{R_{ETC}C}\right)^C \quad (15)$$

For oligomycin treated cells, since ATP synthase in inhibited by oligomycin, we can neglect term $R_{ATP}$. Similarly to control cells, we write

$$\left(\frac{1}{RC}\right)^O = \underbrace{\left(\frac{1}{R_{ATP}C}\right)^O}_{=0} + \left(\frac{1}{R_{Leak}C}\right)^O + \left(\frac{1}{R_{ETC}C}\right)^O \quad (16)$$

Our goal is to calculate $(1/R_{ATP}C)^C$ from equation (15). Since the left-hand terms of equations (14,15) are specified as the mean values from experimental data, we need an expression for $(1/R_{Leak}C)^C$ and $(1/R_{Leak}C)^O$ terms. As the leakage current ($I_{Leak}$) is known to be voltage-dependent with a non-ohmic scaling $I_{Leak} \propto e^{\gamma V}$ [28,32,33], the resistance term $R_{Leak}$ is voltage-dependent as well, given that the capacitance $C$ is assumed constant. Therefore, we write a function that is voltage-dependent $l(V) = (1/R_{Leak}C)$ . Consequently, the term $(1/R_{ATP}C)$ is voltage-dependent and we write $a(V) = (1/R_{ATP}C)$. We assume that that the time constants for electron transport chain are the same in control and oligomycin treated cells $(1/R_{ETC}C)^C = (1/R_{ETC}C)^O = (1/R_{ETC}C)$. In the case of control in equation (16), we can write:

$$\left(\frac{1}{RC}\right)^C = a(V) + l(V) + \left(\frac{1}{R_{ETC}C}\right) \quad (17)$$

However, in case of oligomycin in equation (17) we can treat the term $l(V) = (1/R_{Leak}C)^O$ as approximately constant at a representative median voltage $\bar{V}_o$ (median voltage hyperpolarization with oligomycin), write $l(\bar{V}_o\ ) = (1/R_{Leak}C)^O$and obtain:

$$\left(\frac{1}{RC}\right)^O = l(\bar{V}_o\ ) + \left(\frac{1}{R_{ETC}C}\right) \quad (18)$$

Subtracting equation (18) from equation (17), we obtain:

$$\left(\frac{1}{RC}\right)^{C} - \left(\frac{1}{RC}\right)^{O} = a(V) + l(V) - l(\bar{V}_o) \tag{19}$$

To extract an expression for $a(V)$, we need an expression for leakage term $l(V)$. To achieve this, we use previously reported scaling between leakage current and voltage [33] as follows: $I_{Leak} \propto e^{\gamma V}$, where $\gamma = 0.0122$. From Ohm's law, $I_{Leak} = V/R_{Leak}$, we can write $R_{Leak}C \propto Ve^{-\gamma V}$. Therefore, we express the function $l(V)$ using $l_o$ as a proportionality constant:

$$l(V) = l_o e^{\gamma V}/V \tag{20}$$

Inserting equation (20) into equation (19), we obtain:

$$\left(\frac{1}{RC}\right)^{C} - \left(\frac{1}{RC}\right)^{O} = a(V) + l_o e^{\gamma V}/V - l_o e^{\gamma \bar{V}_o}/\bar{V}_o \tag{21}$$

To specify the proportionality constant $l_o$, we set $V = V_{G2}$, which is the membrane potential in G2 phase (prior to the mitotic mitochondrial hyperpolarization). We then quantified the ATP synthesis to leakage ratio ($I_{ATP}/I_{Leak}$) by measuring the oligomycin sensitive and insensitive proportions of mitochondrial oxygen consumption using a Seahorse Extracellular Flux Analyzer in L1210 cells (Supplementary Figs. 7A, 7C). Based on these measurements we set $I_{ATP}/I_{Leak}|_{V=V_{G2}} = \beta$ where $\beta = 4.88$. Using Ohm's law, we write $a(V_{G2})/l(V_{G2}) = \beta$. We then set $V = V_{G2}$ (-150 mV) in equation (21) and obtain:

$$\left(\frac{1}{RC}\right)^{C} - \left(\frac{1}{RC}\right)^{O} = (1+\beta) l_o e^{\gamma V_{G2}}/V_{G2} - l_o e^{\gamma \bar{V}_o}/\bar{V}_o \tag{22}$$

We can therefore solve equation (22) with respect to $l_o$ and obtain:

$$l_o = \frac{\left(\frac{1}{RC}\right)^C - \left(\frac{1}{RC}\right)^O}{\frac{(1+\beta)e^{\gamma V_{G2}}}{V_{G2}} - \frac{e^{\gamma \bar{V}_o}}{\bar{V}_o}} \tag{23}$$

Inserting equation (23) into equation (19), we obtain an expression for time constant $(1/R_{ATP}C)^C_{CDK1\,off}$. Note that now we have added the subscript 'CDK1 off' for clarity:

$$\begin{aligned}\left(\frac{1}{R_{ATP}C}\right)^C_{CDK1\,off} &= \left(\frac{1}{RC}\right)^C_{CDK1\,off} - \left(\frac{1}{RC}\right)^O_{CDK1\,off} \\ &- \left(\frac{\left(\frac{1}{RC}\right)^C_{CDK1\,off} - \left(\frac{1}{RC}\right)^O_{CDK1\,off}}{\frac{(1+\beta)e^{\gamma V_{G2}}}{V_{G2}} - \frac{e^{\gamma \bar{V}_o}}{\bar{V}_o}}\right)\left(\frac{e^{\gamma V}}{V} - \frac{e^{\gamma \bar{V}_o}}{\bar{V}_o}\right)\end{aligned} \tag{24}$$

***$I_{ATP}$* for CDK1$_{on}$**

In the case of CDK1$_{on}$ we apply the same methodology for control and oligomycin cells as in the case of CDK1$_{off}$, as shown in equations (15-23). We also assume that the inverse time constants in leakage in CDK1$_{on}$ and CDK1$_{off}$ are connected through the relationship of equation (25), where $c$ is a constant:

$$l_{CDK1\,on}(V) = l(V) + c \tag{25}$$

For CDK1$_{on}$ we can therefore write:

$$\left(\frac{1}{R_{ATP}C}\right)^{C}_{CDK1\,on} = \left(\frac{1}{RC}\right)^{C}_{CDK1\,on} - \left(\frac{1}{RC}\right)^{O}_{CDK1\,on} - \left(\frac{\left(\frac{1}{RC}\right)^{C}_{CDK1\,off} - \left(\frac{1}{RC}\right)^{O}_{CDK1\,off}}{\frac{(1+\beta)e^{\gamma V_{G2}}}{V_{G2}} - \frac{e^{\gamma \bar{V}_o}}{\bar{V}_o}}\right)\left(\frac{e^{\gamma V}}{V} - \frac{e^{\gamma \bar{V}_o}}{\bar{V}_o}\right) \tag{26}$$

Having specified the values for $R_{ATP}C$ for both CDK1 on and off, we can calculate the ATP currents using equations (13,14). For the $\Delta t$ regions, where CDK1 transitions from off to on ($t = [t_0, t_1]$) and vice versa ($t = [t_2, t_3]$), we assumed linear interpolations between the ATP currents at the transition points.

To extract mean and standard deviation of ATP currents ($I_{ATP}$) for each cell, we utilized the population average $R_{ATP}C$ values and the TMRE measurement-based estimates of voltage. We first linearly interpolated the current of each cell obtained from equations (13) and (14) to have 1,000 data points around mitosis (starting from 2 h before and ending at 1.5 h after metaphase-anaphase transition ($t = t_f$)). Then, we aligned each interpolated current trace at the metaphase-anaphase transition and calculated mean ± standard deviation at each time point that was linearly interpolated.

##### *General implications of RC values*

Our calculations revealed that $R_{ATP}C$ is higher during $CDK1\ on$ state than during $CDK1\ off$ state, $0.7 \pm 0.2$ and $0.27 \pm 0.01$ h respectively (mean ± s.e.m.). These values are the mean $R_{ATP}C$ values over a voltage range of -150 mV to -200mV (Supplementary Fig. 11C). Our data also revealed that in control cells $RC_{CDK1\,on}/RC_{CDK1\,off} = 1.4 \pm 0.1$ (mean ± s.e.m.), which is statistically different from the value 1.0 ($p = 1.8 * 10^{-4}$, Student's unpaired t-test, n=85). Thus, the overall time constant $RC$ increases in CDK1 $on$ state. Note that the overall time constant $RC$ is determined, in addition to $R_{ATP}C$ , by the two other time constants $R_{ETC}C$ and $R_{Leak}C$ based on equation (15). We therefore questioned if $R_{ETC}C$ and/or $R_{Leak}C$ may also change during CDK1 $on$ state. To test this hypothesis, we assumed that $R_{ETC}C$ and $R_{Leak}C$ stay constant between $CDK1\ on$

and $off$ states, with only $R_{ATP}C$ becoming higher during $CDK1$ $on$ state, as shown above. This hypothetical ratio $RC_{CDK1\ on}/RC_{CDK1\ off} = 1.8 \pm 0.7$ is significantly different from our measured ratio of $1.4 \pm 0.1$ (mean ± s.e.m.) ($p = 8 * 10^{-4}$, Student's unpaired t-test, n=85). Therefore, our model predicts that $R_{ETC}C$ and/or $R_{Leak}C$ decrease during $CDK1$ $on$.

This prediction of our model that $R_{ETC}C$ and/or $R_{Leak}C$ decrease (i.e. ETC activity and/or leakage increase) during mitosis ($CDK1$ $on$ state) is consistent with literature. It has been reported that ETC activity increases during mitosis [39-41], and our observation that TMRE increases upon mitotic entry in oligomycin treated cells also implies increased ETC activity during mitosis (Figs. 3A, 3B). However, when we synchronized cells to mitosis and measured their oxygen consumption rate in comparison to G2 arrested cells, we did not observe increased oxygen consumption (Supplementary Figs. 7D, 7E). This likely reflects an artefact of cell cycle synchronization, as mitotic arrests are known to result in inhibition of cell growth and degradation of mitochondria [3,40]. In addition, the mitotically arrested cells displayed higher mitochondrial leakage than G2 arrested cells (Supplementary Figs. 7D, 7E), implying that $R_{Leak}C$ is also decreased during mitosis. As leakage also increases with increased ΔΨm [33], mitotic cells should display increased mitochondrial proton leakage. Mitochondrial proton leakage is the main heat source within cells [42,43] and, consistently, cells in early mitosis have been shown to produce more heat [44]. Thus, while our model cannot quantify the extent to which $R_{ETC}C$ and $R_{Leak}C$ change, it seems likely that both values decrease during mitosis ($RC_{CDK1\ on}$ state) and thus mitotic cells would display increased $I_{ETC}$ and $I_{Leak}$.

#### Supplementary Note 8. Calculating ratios and integral of current $I_{ATP}/C$

As the key conclusion of our model, we calculate the extent of ATP synthesis rate change between G2 and mitotic cells, which we call $r_{I_{ATP}}$. To do this, we obtain the ratio between the currents $I_{ATP}/C$ from equations (13,14):

$$r_{I_{ATP}} = \frac{I_{ATP,CDK1\,on}}{I_{ATP,CDK1\,off}} \tag{27}$$

To calculate $r_{I_{ATP}}$ from equation (27), we used the minimum value of mean $I_{ATP,CDK1\,on}$ and the baseline (G2) value of mean $I_{ATP,CDK1\,off}$ of control cells.

We also integrated the equations (13,14) using the trapezoidal rule method over discrete points ($t_i$ , $I_{ATP}/C_i$ ) to obtain the total charge transported across the inner mitochondrial membrane through ATP synthase, i.e. the amount of ATP synthesized. Here, $I_{ATP}/C_i$ can refer either to $CDK1_{on}$ or $CDK1_{off}$:

$$Q_{I_{ATP}} = \sum_{i=2}^{N} \frac{I_{ATP}/C_i + I_{ATP}/C_{i-1}}{2} \Delta t = \left( \frac{I_{ATP}/C_0}{2} + \sum_{i=2}^{N-1} I_{ATP}/C_i + \frac{I_{ATP}/C_N}{2} \right) \Delta t \tag{28}$$

Note that in equation (28) we did the integration over the mean ATP synthesis rate we derived from control sets, which we had interpolated (Supplementary Note 7). Due to the interpolation in time $t$, the time step was constant, $\Delta t = t_{i+1} - t_i = constant$.

The integration was done for ATP currents in early mitosis (spanning 30 min from G2/M transition to metaphase-anaphase transition) and for anaphase (spanning 15 minutes from metaphase-anaphase transition to the approximate end of anaphase) [3,5]. As a reference, these values were compared to a hypothetical situation where the ATP synthesis rate remained at G2 levels.

#### Supplementary Note 9. Error analysis

We calculate the error in time constant $(R_{ATP}C)$ of equations (24 and 26) for CDK1 on and off respectively, by accounting for the error in the estimation of time constants $(RC)^C$, $(RC)^O$, where superscript C refers to control and O to oligomycin data. We did not include error of voltage $V$ in these calculations as the extent to which our TMRE-to-voltage conversion factors and our voltage dependent leakage factors influence the voltage values is not known and thus we cannot directly estimate their contribution to the error. Instead, we carried out sensitivity analyses on the TMRE-to-voltage conversion (Supplementary Fig. 12) and the voltage-dependent leakage (Supplementary Fig. 14, Supplementary note 10). Note also that our the TMRE measurements of mitotic cells is typically below 2 % (Supplementary Fig. 1). Below, we do the error analysis for CDK1 on and off separately.

***Case CDK1$_{off}$***

To estimate the error $s_f$ of a function $f$ which depends on two variables $x, y$ with errors $s_x, s_y$ we use equation (29), assuming that $x, y$ are independent, uncorrelated, variables:

$$s_f = \sqrt{\left(\frac{\partial f}{\partial x} s_x\right)^2 + \left(\frac{\partial f}{\partial y} s_y\right)^2} \tag{29}$$

For simplicity in writing of equation (24) we can use the following symbols, where $\delta(RC)^C_{CDK1\,off}$ and $\delta(RC)^O_{CDK1\,off}$ stand for standard errors for $(RC)^C_{CDK1\,off}$ and $(RC)^O_{CDK1\,off}$ respectively:

$$f_{off} = (R_{ATP}C)^C_{CDK1\,off} \tag{30}$$

$$x = (RC)^C_{CDK1\,off} \tag{31}$$

$$y = (RC)^O_{CDK1\,off} \tag{32}$$

$$s_x = \delta(RC)^C_{CDK1\,off} \tag{33}$$

$$s_y = \delta(RC)^O_{CDK1\,off} \tag{34}$$

Note that from our datasets we obtained $s_x/x = 4\%$ and $s_y/y = 8\%$ (Fig. 4D), which means that the standard errors are relatively small with respect to the mean values, thus justifying the use of equation (29).

As stated above, we assume that the rest of the terms carry no error. Thus, we can group the terms together under a variable G:

$$G = \left( \frac{1}{\frac{(1+\beta)e^{\gamma V_{G2}}}{V_{G2}} - \frac{e^{\gamma \bar{V}_o}}{\bar{V}_o}} \right) \left( \frac{e^{\gamma V}}{V} - \frac{e^{\gamma \bar{V}_o}}{\bar{V}_o} \right) \tag{35}$$

Therefore equation (24) can be written as:

$$f_{off} = \frac{1}{\frac{1-G}{x} + \frac{G-1}{y}} \tag{36}$$

We take the derivative of equation (34) with respect to $x$ and $y$ and write this in compact form since each derivative is proportional to $f_{off}^2$:

$$\frac{\partial f_{off}}{\partial x} = -\frac{(1-G) f_{off}^2}{x^2} \tag{37}$$

$$\frac{\partial f_{off}}{\partial y} = -\frac{(G-1) f_{off}^2}{y^2} \tag{38}$$

Therefore, we estimate the error $\delta(R_{ATP}C)^C_{CDK1\,off}$ combining equations (29-38):

$$\delta(R_{ATP}C)^C_{CDK1\,off} = \left[(R_{ATP}C)^C_{CDK1\,off}\right]^2 \mathrm{x}$$

$$|\quad -1| \sqrt{\left( \frac{\delta(RC)^C_{CDK1\,off}}{[(RC)^C_{CDK1\,off}]^2} \right)^2 + \left( \frac{\delta(RC)^O_{CDK1\,off}}{[(RC)^O_{CDK1\,off}]^2} \right)^2} \tag{39}$$

From equation (39) it is clear that the error $\delta(R_{ATP}C)^{C}_{CDK1\,off}$ depends on voltage $V$ included in the expression of $G$ from equation (35).

***Case CDK1$_{on}$***

For simplicity with symbols, we set:

$$f_{on} = (R_{ATP}C)^{C}_{CDK1\,on} \tag{40}$$

$$z = (RC)^{C}_{CDK1\,on} \tag{41}$$

$$w = (RC)^{O}_{CDK1\,on} \tag{42}$$

$$s_z = \delta(RC)^{C}_{CDK1\,on} \tag{43}$$

$$s_w = \delta(RC)^{O}_{CDK1\,on} \tag{44}$$

Note that in this case we also have $s_z/z = 8\%$ and $s_w/y = 12\%$ (Fig. 4D) which means that the standard errors are relatively small with respect to the mean values, thus justifying the use of an equation similar to (29) but with four terms.

$$s_f = \sqrt{\left(\frac{\partial f}{\partial x}s_x\right)^2 + \left(\frac{\partial f}{\partial y}s_y\right)^2 + \left(\frac{\partial f}{\partial z}s_z\right)^2 + \left(\frac{\partial f}{\partial w}s_w\right)^2} \tag{45}$$

Similar to the case CDK1$_{on}$, we apply equations (31-34, 40-45) to equation (26) and obtain:

$$f_{on} = \frac{1}{\frac{1}{z} - \frac{1}{w} - \frac{G}{x} + \frac{G}{y}} \tag{46}$$

We take the derivatives of equation (46) with respect to $z, w, x, y$ and write this in compact form since each derivative is proportional to $f_{on}^2$:

$$\frac{\partial f_{on}}{\partial x} = \frac{G f_{on}^2}{x^2} \tag{47}$$

$$\frac{\partial f_{on}}{\partial y} = -\frac{G f_{on}^2}{y^2} \tag{48}$$

$$\frac{\partial f_{on}}{\partial z} = -\frac{f_{on}^2}{z^2} \tag{49}$$

$$\frac{\partial f_{on}}{\partial w} = \frac{f_{on}^2}{w^2} \tag{50}$$

Therefore, we estimate the error $\delta(R_{ATP}C)^{C}_{CDK1\,on}$ by combining equations (40-50):

$$\delta(R_{ATP}C)^{C}_{CDK1\,on} = [(R_{ATP}C)^{C}_{CDK1\,on}]^2 \text{ x}$$

$$\sqrt{\left(\frac{G\delta(RC)^{C}_{CDK1\,off}}{[(RC)^{C}_{CDK1\,off}]^2}\right)^2 + \left(\frac{G\delta(RC)^{O}_{CDK1\,off}}{[(RC)^{O}_{CDK1\,off}]^2}\right)^2 + \left(\frac{\delta(RC)^{C}_{CDK1\,on}}{[(RC)^{C}_{CDK1\,on}]^2}\right)^2 + \left(\frac{\delta(RC)^{O}_{CDK1\,[illegible]}}{[(RC)^{O}_{CDK1\,on}[illegible]}\right.} \tag{51}$$

Similar to equation (39), the error $\delta(R_{ATP}C)^{C}_{CDK1\,on}$ depends on voltage $V$ included in the expression of $G$ from equation (35).

***Calculating errors for currents $I_{ATP}$/C, its ratios and integrals***

Using expressions (39, 51) for the errors for $(R_{ATP}C)^C$ we calculate the error in the current $I_{ATP}/C$ from equations (13,14). Applying the same methodology we used for deriving equations (39,51), we obtain:

$$\delta\left(\frac{I_{ATP,CDK1\,off}}{C}\right) = \frac{I_{ATP,CDK1\,off}}{C}\left|\frac{\delta(R_{ATP}C)^{C}_{CDK1\,off}}{(R_{ATP}C)^{C}_{CDK1\,off}}\right| \tag{52}$$

$$\delta\left(\frac{I_{ATP,CDK1\,on}}{C}\right) = \frac{I_{ATP,CDK1\,on}}{C}\left|\frac{\delta(R_{ATP}C)^{C}_{CDK1\,on}}{(R_{ATP}C)^{C}_{CDK1\,on}}\right| \tag{53}$$

We calculate the error for $r_{I_{ATP}}$ in a similar fashion as before:

$$\delta r_{I_{ATP}} = \frac{R_{ATP,CDK1\,off}}{R_{ATP,CDK1\,on}} \sqrt{\left(\frac{\delta (R_{ATP}C)^{C}_{CDK1\,off}}{(R_{ATP}C)^{C}_{CDK1\,off}}\right)^2 + \left(\frac{\delta (R_{ATP}C)^{C}_{CDK1\,on}}{(R_{ATP}C)^{C}_{CDK1\,on}}\right)^2} \tag{54}$$

Using a formula of partial derivatives for the errors similar to equations (29,45), we calculate the error of the integral, $\delta Q_{I_{ATP}}$ as:

$$\delta Q_{I_{ATP}} = \Delta t \sqrt{\left[\delta\left(\frac{I_{ATP}/C_0}{2}\right)\right]^2 + \sum_{i=2}^{N-1} [\delta(I_{ATP}/C_i)]^2 + \left[\delta\left(\frac{I_{ATP}/C_N}{2}\right)\right]^2} \tag{55}$$

#### Supplementary Note 10. Sensitivity analysis for model

First, we carried out a sensitivity analysis on the selection criterion for the best $Split$ value (Supplementary Fig. 10E). In addition to selecting the $Split$ value based on maximization of R-squared, we also selected the $Split$ value based on maximization of $RC$ value confidence (Supplementary Fig. 11B). We defined the $RC$ value confidence as $|RC_{max} - RC_{min}|/RC$, where $RC_{max}, RC_{min}$ represent the maximum and minimum value of $RC$ with 95% confidence. We observed little difference between the two selection criteria (Supplementary Fig. 11B, mean values of $RC$) and thus decided to use the maximization of R-squared as a selection criterion for the $Split$ value.

After establishing the maximization of R-squared as a selection criterion, we carried out sensitivity analyses on several of the values that our modeling assumes or that we obtain from literature. In these analyses we focused on a single main prediction made by our model: the ratio of ATP synthesis between mitosis and G2 ($\frac{I_{ATP,CDK1\,on}}{I_{ATP,CDK1\,off}}$). Our reference value $r_{I_{ATP}} = \frac{I_{ATP,CDK1\,on}}{I_{ATP,CDK1\,off}} = 46\ \% \pm 11\ \%$ (mean ± s.e.m.), and was based on the following assumptions:

1) Baseline voltage $V_o = -150\ mV$ (Supplementary note 6)
2) TMRE-to-ΔΨm conversion factor $c_2 = 0.0215$ (Supplementary note 6)
3) For ΔΨm-dependent leakage $\beta = 4.88, \gamma = 0.0122$ (Supplementary note 7)
4) Start angle for data fitting ($\theta_{start} = 15\ degrees$) (Supplementary note 6, Supplementary Fig. 10B)
5) The duration of CDK1 turning on and off ($\Delta t = 8.57\ min$) (Supplementary note 6, Supplementary Fig. 6).

First, we calculated $r_{I_{ATP}}$ when changing the parameters of TMRE-to-ΔΨm conversion factor ($c_2 = 0.0185, 0.02, 0.0215, 0.0230, 0.0245$) and the assumed baseline voltage ($\Delta\Psi m = -140, -151, \dots, -160\ mV$) while keeping the rest of the parameters fixed. Over the range of these parameters $r_{I_{ATP}}$ was fluctuating within the range $35 - 50\%$ (Supplementary Fig. 12A), indicating that our model is not sensitive to the values used for TMRE-to-voltage conversion. We also calculated the relative error (Supplementary Fig. 12B) in the estimation of $r_{I_{ATP}}$ based on the equation:

$$R.E.\delta r_{I_{ATP}} = \frac{\delta r_{I_{ATP}}}{r_{I_{ATP}}} \tag{56}$$

We observed that $R.E.\delta r_{I_{ATP}}$ is kept below 30% (Supplementary Fig. 12C) throughout this range. In other words, the extent of our uncertainty in the estimation of $r_{I_{ATP}}$ is relatively small. Moreover, we calculated the data inclusion based on our criteria, as well as fitting quality (R-squared of the fits), and observed that throughout the range, over 95% of data were preserved (Supplementary Fig. 12B) and the mean quality of the fits stays consistently high (average $R^2 > 0.94$) (Supplementary Fig. 12D).

Second, we calculated $r_{I_{ATP}}$ when changing the parameters used to define fitting regions. We changed the parameters used for to define where fitting starts ($\theta_{start} = 5{,}10{,}15{,}20\ degrees$) and the duration of CDK1 on/off transitions ($\Delta t = 6{,}7{,}8.57{,}9{,}10\ min$), while keeping the rest of the parameters fixed. Over the range of these parameters $r_{I_{ATP}}$ fluctuated within the range $15-50\%$ (Supplementary Fig. 13A). The $r_{I_{ATP}}$ change was more pronounced when $\theta_{start}$ changed. However, we note that a lower starting angle ($\theta_{start} = 5-10\ degrees$) is associated with more noise in the data leading to a higher $R.E.\delta r_{I_{ATP}}$ (Supplementary Fig. 13C). Notably, the $R.E.\delta r_{I_{ATP}}$ was minimized at our reference values. In addition, more single-cell data sets were excluded when $\theta_{start}$ and $\Delta t$ decreased (Supplementary Fig. 13B) (see Supplementary note 6 for data exclusion criteria), although the R-squared of fits remained high (Supplementary Fig. 13D).

Third, we calculated $r_{I_{ATP}}$ for multiple pairs of the values $(\beta, \gamma)$, which define the voltage-dependent leakage in our model. For each pair from combining values ($\beta = 4, 4.88, 6$) and ($\gamma = 0, 0.005, 0.0122{,}0.015, 0.02, 0.03$), we computed $r_{I_{ATP}}$ over a sweep of the parameters for TMRE-to-ΔΨm conversion factor ($c_2 = 0.0185, 0.02{,}0.0215, 0.0230, 0.0245$) and baseline voltage ($\Delta\Psi m = 140, 151, \ldots, 160$). Over these ranges, we observed that $r_{I_{ATP}}$ fluctuated within the range $35-55\%$ (Supplementary Fig. 14). Note that at $\gamma = 0$ the leakage current has an ohmic scaling with ΔΨm. Thus, our sensitivity analysis overall indicates that the results of our model have little sensitivity to the assumed and/or literature-derived values. We conclude that the result of our model, i.e. that L1210 cells decrease their mitochondrial ATP synthesis approximately 55% during early mitosis when compared to G2 levels, is reflecting the underlying biological changes in mitosis and not a modeling artefact.

#### Materials and Methods

##### Cells and cell culture conditions

L1210, BaF3, DT40, Fl5.12 and primary T cells were cultured in RPMI that contained 10 % FBS (Gibco), 11 mM glucose, 2 mM glutamine, 1 mM Na pyruvate, 20 mM HEPES and antibiotic/antimycotic. In addition, Fl5.12 culture media was supplemented with 10 ng/ml IL-3 (R&D Systems), DT40 culture media was supplemented with 3 % chicken serum (Sigma-Aldrich) and primary T cell culture media was supplemented with 10 mM 2-mercaptoethanol, 100 U/ml IL-2 (R&D Systems) and 2 mg/ml anti-human CD28 (BioLegend). S-HeLa cells were grown in DMEM that was supplemented with 10 % FBS, 1 mM sodium pyruvate and antibiotic/antimycotic. L1210 cells were also grown in modified RPMI when comparing growth conditions (Supplementary Figs. 3B, 3C). For these experiments, we used RPMI containing 4 % FBS and either high (25 mM) glucose, low (4 mM) glucose or 10 mM galactose with no glucose.

Isolation of naïve CD3+ and CD8+ primary T cells was carried out from unpurified buffy coat (Research Blood Components) from which PBMCs were isolated using Ficoll-Paque Plus density gradient (GE). After isolation of the PBMCs, the cells were subjected to red blood cell lysis using ACK lysis buffer (Thermo Fisher Scientific). The PBMCs were then washed three times and T cells were isolated using Naive CD3+ or CD8+ T Cell Isolation Kit (Miltenyi Biotec) according to kit instructions. The cells were then activated by culturing the cells on an anti-CD3 coated cell culture plate in the media detailed above. T cells were used for experiments approximately 30 h after activation. The activation of T cells was validated by monitoring cell volumes and counts using coulter counter (Beckman Coulter).

Within all SMR experiments, the culture media used was identical to that described above, apart from any indicated fluorescent probe and/or chemical inhibitor addition. All cell culture reagents were obtained from Invitrogen, unless otherwise stated. L1210 cells we obtained from ATCC (Cat# CCL-219), BaF3 cells were obtained from RIKEN BioResource Center (Cat# RCB4476), DT40 cells, which harbor a CDK1as [45], were a gracious gift from K. Samejima and B. Earnshaw from University of Edinburgh, Fl5.12 cells were a gracious gift from M. Vander Heiden from Massachusetts Institute of Technology, and S-HeLa cells were a gracious gift from K. Elias from Brigham Women's Hospital. All cell lines were tested to be free of mycoplasma.

#### Generation of reporter cell lines

The FUCCI cell cycle marker expressing L1210 cells were generated in a previous study [2]. L1210 and BaF3 cells that stably express an ATP reporter (A-Team), reactive oxygen species reporter (roGFP2-Orp1) or glutathione redox potential reporter (Grx1-roGFP2) were generated using lentiviral vectors obtained from AddGene [19-22]. pEIGW roGFP2-Orp1 was a gift from Tobias Dick (Addgene plasmid #64993 ; http://n2t.net/addgene:64993 ; RRID:Addgene_64993), pEIGW Grx1-roGFP2 was a gift from Tobias Dick (Addgene plasmid # 64990 ; http://n2t.net/addgene:64990 ; RRID:Addgene_64990), ATeam1.03-nD/nA/pcDNA3 was a gift from Takeharu Nagai (Addgene plasmid # 51958 ; http://n2t.net/addgene:51958 ; RRID:Addgene_51958). Lentiviruses for the genetically encoded sensors were produced by calcium phosphate transfection of plasmids into HEK-293T cells. For each sensor, the corresponding lentiviral vector was co-transfected with separate plasmids expressing VSV-G and Gag/Pol. Following a media change after an overnight transfection, virus aliquots were harvested at 48 h and 72 h post-transfection and filtered through 0.45 μm filters. Lentiviruses for containing Ca2+ reporter (GCaMP3) [23] vectors were obtained from Kerafast (cat# FCT188). Note that all genetic sensors were expressed under strong promoters, but FUCCI cell cycle reporter was expressed under the endogenous Geminin promoter.

Lentiviruses were transfected into L1210 and BaF3 cells using spinoculation as described previously [3]. Briefly, approximately $1.5x10^5$ cells were mixed with 10 mg/ml polybrene (EMD Millipore) and the lentiviruses. This mixture was centrifuged at 800 g for 60 min at 25°C, after which the cells were moved to normal culture media for 12 h. This was repeated 4 times, and 24 h later selection process was started with puromycin or neomycin, depending on the transfected vector. Following 5 days of selection, a subpopulation with high fluorescence reporter expression was sorted out using BD FACS Aria.

#### Suspended microchannel resonator (SMR) setup

The SMR devices were fabricated as previously described [46] and carried out at CEA-LETI, France. Exact dimension and geometry of the device can be found in [5]. The SMR devices were actuated the second vibration mode by a piezo-ceramic placed underneath the chip. The SMR were operated in closed-loop, where the output motion of the cantilever is amplified, delayed and fed back to the piezo-ceramic to drive the cantilever. The signal delay was chosen so that the amplitude of the cantilever motion was maximized. The amplification was set to minimize the resonant frequency error, but not in the limit where it distorts the linearization of the resonant frequency. A digital platform previously described [47] was implemented to track the change in resonant frequency (frequency of the closed-loop signal) over time. This close-loop operation configuration results in measurement bandwidth ~1500 Hz, wide enough to capture the frequency modulation during cell transit event, which is typically set to ~150-200 ms. The resonance frequency changes were analyzed and converted to buoyant mass as detailed in [5]. The serial SMR experiments (Fig. 3E) were carried out as detailed previously [3,46,48].

**Optical detection setup for SMR**

The optical setup used in this study is similar to our previous setup [5], except few additions/modifications in optical components to achieve simultaneous measurements of TMRE and FUCCI signals. First, we implemented a "full multiband configuration", which is composed of a multiband exciter filter (Semrock, FF01- 387/485/559/649-25), a multiband emission filter (Semrock, FF01- 440/521/607/694/809-25) and a multiband dichroic beam splitter (Semrock, FF 408/504/581/667/762-Di01-25x36). This filter set cube was mounted in our microscope (Nikon). Second, a dichroic beam splitter (Semrock, FF580-FDi01-25x36) was mounted beyond the image plane to separate the multi-color fluorescence output signal into single-color fluorescent signals (i.e., FUCCI and TMRE signals). The split fluorescent signals were measured by photomultiplier tubes (Hamamatsu, H10722-20) mounted on top of the microscope with single band pass filters (Semrock, FF01-520/35 for FUCCI, Semrock, FF01-607/35 for TMRE, respectively). Third, to minimize phototoxicity from long-term measurements, the excitation light source was kept on for less than 500 ms for each measurement, during which the cell was flown through the excitation light path. The cell was directly exposed to the light for approximately 50 ms during each measurement. This fluorescence measurement was carried out approximately every 2 min (in conjugation with every second buoyant mass measurement) (Supplementary Figs. 1A, 1B). The area of light exposure was limited 60X60 μm and the area of emission collection was limited to 40X60 μm. We did not observe changes to single-cell growth rates when measuring TMRE signal with this setup. All parts of the optical detection setup (including objective lens: 50×/0.55-NA Nikon-CFI, LU Plan ELWD WD 10.1 mm; rectangular slit for emission area control: Thorlab;

light source: Lumencor, Spectra X Light Engine) were identical to those used before [5]. The optical path for simultaneous two-color measurements is similar to what has been shown in the previous work [2].

**System operation and error quantification**

Buoyant mass and fluorescence signal(s) (TMRE alone or TMRE and FUCCI) were continuously measured using previously explained hydrodynamic trapping approach [2,3,5]. The hydrodynamic trapping, SMR measurements and optical operation is explained in more detail in (Supplementary Figs. 1A, 1B).

We quantified the error in optical measurement by repeatedly measuring fluorescently labeled fixed cells (labelling detailed below). We then carried out linear fitting to the data and calculated the average deviation of each measurement from a linear fitting. The signal-to-noise ratio displayed in (Supplementary Figs. 1C, 1D) was calculated by dividing the mean fluorescence intensity of each cell by the standard deviation of the measurements from the fitted line. Using this approach, we estimated our optical measurements to have an error (CV) of approximately 2% for mitotic cells. See (Supplementary Figs. 1C, 1D) for details. The error in buoyant mass measurement were quantified previously (by repeatedly measuring a 10 μm polystyrene bead) to be <0.1 pg, which corresponds to an error of <0.25% [3].

**Detecting cell cycle transitions within the SMR**

Three cell cycle transitions were assigned for each cell in the SMR data. First, the G2/M transition was detected using node deviation signal (Supplementary Fig. 5), which is an acoustic, stiffness dependent signal detected by the SMR [5]. This timing was validated also by measurements of mitotic cell swelling (Fig. 1E), which start immediately following mitotic entry [1,13]. Single-cell density was measured as detailed in [1]. Second, the metaphase-anaphase transition was detected using the node deviation signal (Supplementary Fig. 5) and also using the rapid change in buoyant mass accumulation rates [3]. The timing of metaphase-anaphase transition was further validated using the FUCCI cell cycle sensor expressing cells (Figs. 1F, 1G). The imaging of FUCCI cells (Fig. 1F) was carried out using IncuCyte. Third, the abscission of the daughter cells was assigned using the sudden ~50 % reduction in buoyant mass (Fig. 1A). In addition, we assigned anaphase to last 15 min (Fig. 4E), as based on our previous quantifications of L1210 cell elongation duration [5].

**Membrane potential and mitochondrial content measurements**

For all cells, mitochondrial membrane potential was examined using the fast equilibrating small-molecule probe Tetramethylrhodamine ethyl ester perchlorate (TMRE) (Invitrogen) in 10 nM (non-quenching) concentration, with the exception that DT40 cells and primary T cells were measured using 20 nM TMRE. TMRE was detected on the TRICT channel of our SMR setup (see above). For a typical experiment, L1210 cells at an approximate confluency of 300.000 cells/ml were stained with 10 nM TMRE for 30 min, after which cells were loaded in to the SMR. Within the SMR, cells were grown in the presence of 10 nM TMRE. Mitochondrial membrane potential was also measured using 10 µM (quenching) concentrations of the Rhod123 (Invitrogen). Rhod123 was detected on the FITC channel of our SMR setup (see above). Cells were stained for 45 min with 10 µM Rhod123, washed twice with media and loaded in to the SMR. No Rhod123 was present in the media within the SMR. Note that Rho123 slowly diffuses out of the cell and this excludes long term experiments and reduces the quantitative accuracy of prolonged analyses. In order to observe mitosis before Rhod123 diffused out of the cell, we carried out Rho123 experiments by loading only large G2 cells in to the SMR. For both TMRE and Rhod123, the quenching and non-quenching states were validated by staining L1210 cells as described above and quantifying fluorescence levels after 30 min treatment with 1 µM FCCP or oligomycin using BD Biosciences flow cytometer LSR II HTS with excitation lasers at 488 nm and 561 nm, and emission filters at 530/30 and 585/15.

When analyzing the error of mitochondrial fluorescence measurements, L1210 cells were stained for 30 min with MitoTracker Red CMXRos (Invitrogen), after which cells were washed with PBS and fixed in 4 % PFA for 10 min. The cells were then washed, and the mitochondrial signal of each cell was repeatedly measured in SMR using the TRICT channel of our SMR setup (see above).

Mitochondrial content was examined using MitoTracker Green probe (Invitrogen) in 50 nM concentration. MitoTracker Green was detected on the FITC channel of our SMR setup (see above). As with TMRE, cells were stained for 30 min after which cells were loaded in to the

SMR. MitoTracker Green was present in the culture media within SMR. Note that in these concentrations the presence of TMRE and MitoTracker Green did not affect cell growth rate.

Plasma membrane potential was examined using Bis-(1,3-dibutylbarbituric acid) trimethine oxonol ($DiBAC_4(3)$) probe (Invitrogen) in 1 μM concentration. $DiBAC_4(3)$ was detected on the FITC channel of our SMR setup (see above). Cells were stained for 45 min, after which cells were loaded in to the SMR. Within the SMR, cells were grown in the presence of $DiBAC_4(3)$.

When using flow cytometer to measure live cell mitochondrial or plasma membrane potential, TMRE and $DiBAC_4(3)$ were used at 10 nM and 1 μM concentrations, respectively, and cells were analyzed in the presence of the fluorescence probes.

**Chemical perturbations**

Chemical treatments within the SMR were carried out by adding the chemical of interest to the culture media within the SMR. Thus, cells were exposed to the chemical when loading the cells in to SMR, which in a typical experiment was approximately 2 h prior to mitosis. Where separately indicated (Figs. 3D, 3E), chemicals were injected to the SMR culture media during the experiment. All SMR experiments with chemical treatments were stopped following mitotic exit, so that each replicate with chemical inhibitors represents a completely independent experiment.

Unless otherwise indicated, chemical perturbations were done using the following chemical concentrations: 1 μM oligomycin, 1 μM FCCP, 1 μM TPB (Sigma-Aldrich), 5 μM EIPA (Sigma-Aldrich), 1 μg/ml Nocodazole (Sigma-Aldrich), 5 μM STLC (Sigma-Aldrich), 15 μM proTAME, 1 μM RO-3306, 400 nM BMS-265246, 100 nM okadaic acid, 2 μM rotenone, 2 μM antimycin A, 2 mM thymidine. All chemicals were diluted in DMSO, except okadaic acid, which was diluted in EtOH. Unless otherwise indicated, all chemical inhibitors were obtained from Cayman Chemicals. For cell cycle targeting chemicals, treatment efficiency was validated by examining cell cycle profile.

**Cell cycle synchronizations**

In L1210 and BaF3 cells, the cell cycle was synchronized to G2 using double thymidine block followed by G2 arrest as detailed in [3]. Briefly, cells were treated with 2 mM thymidine for 15 h, washed twice with PBS, cultured for 6 h in normal culture media, retreated with 2 mM thymidine for 6 h, washed twice with PBS, returned to normal culture conditions for 3 h and treated with 5 μM RO-3306 for 7 h. At this point approximately 80-90 % of cells were arrested in G2 and upon removal of RO-3306 part of the cells enter mitosis immediately [3].

The cell cycle status of L1210 cells was analyzed as follows: The cells were washed with PBS, fixed in 4 % PFA for 10 min, washed with PBS, permeabilized with 0.5 % Triton X-100 for 10 min, washed with PBS and blocked with 5 % BSA in PBS for 30 min. The cells were then stained with p-Histone H3 (S10) antibody (D2C8, conjugated to Alexa 488, Cell Signaling Technology, #3465S) in a PBS solution containing 5 % BSA o/n at +4°C. The p-Histone H3 antibody was used in the concentration recommended by the supplier. The following day the cells were washed with PBS and stained with 1:2000 dilution of NuclearMask Blue (#H10325, Thermo Fisher Scientific) for 30 min in RT. Finally, the cells were washed three times with PBS, mixed in to PBS supplemented with 1% BSA and put on ice until FACS analysis. The relative proportions of mitotic (4N DNA content, p-Histone positive), G2 cells (4N DNA content, p-Histone negative) and G1&S cell (>4N DNA content, p-Histone negative) was quantified using BD Biosciences flow cytometer LSR II HTS with excitation lasers at 355 nm, 488 nm and 561 nm, and emission filters at 450/50, 530/30 and 585/15.

For quantification of cell cycle status in BaF3 and DT40 cells we used propidium iodine (PI) staining of DNA. The cells were washed with PBS, fixed in 70 % ice cold EtOH o/n, washed with PBS and stained with FxCycle PI/RNase Staining Solution (Invitrogen). After a 30 min staining DNA content was quantified using BD Biosciences flow cytometer LSR II HTS.

**Oxygen consumption measurements**

Oxygen consumption was measured using the XF24 Seahorse Extracellular Flux analyzer. The Seahorse cell culture plates were coated with poly-L-lysine and approximately 80,000 L1210 cells were loaded in to each well. The experiment was started 3 hours after plating the cells. The oxygen consumption measurements were carried out in normal cell culture media (see above) to maximize comparability to the conditions within SMR. Oxygen consumption was measured four times with 6 min intervals, after which cells were treated with 1 μM oligomycin and, four measurements later, with 2 μM rotenone and 2 μM antimycin A. After the experiment, cells in each well were counted and this cell count was used to normalize the data.

When measuring oxygen consumption in cell arrested in G2 and mitosis, G2 arrested cells were washed and moved back in to 5 µM RO-3306 containing media (to maintain G2 arrest) or in to 5 µM STLC containing media (to achieve prometaphase arrest) [3]. The cells were then plated on to the Seahorse cell culture plates and two hours later the oxygen consumption measurement was carried out as described above. During the oxygen consumption measurements, parallel cultures were processed for FACS based cell cycle analysis as described above. The relative proportions of G2 and mitotic cell in the G2 arrested and mitotically arrested populations were used to normalize the oxygen consumption data so that incomplete cell cycle synchrony between G2 and mitosis does not bias the data.

**Data analysis**

All details of the electrical circuit model and data analysis for the model can be found in (Supplementary notes 4-9). The modeling and SMR & fluorescence data processing were carried out using custom MATLAB codes. Quantifications of TMRE increase during mitosis was carried out by normalizing the TMRE signal of each data point to the median signal a 3 h time period prior to daughter cell abscission, after which the highest TMRE value during mitosis was used of quantification.

**Statistics**

In population level experiments, such as cell cycle analyses or oxygen consumption measurements, each replicate represents an independent culture. In control SMR experiments, cell progenies were monitored for various durations, so that each independent experiment can have one or more mitotic events. In contrasts, all chemical treatments within the SMR were carried out so that only one mitotic event was analyzed for each experiment. Details of replicate numbers can be found in figures and figure legends. All experiments were repeated at least three times, and the experiments used for modeling were repeated over 30 times. Importantly, as some data were excluded from the modeling, as cells did not reach a steady TMRE baseline prior to cell division. For full details of data exclusion, please see (Supplementary note 6). Statistical significances are indicated in each figure and the statistical details can be found in the figure legends. For statistical analysis of the electrical circuit model results, please see Supplementary note 9. All statistical analyses were carried out using MATLAB or Origin Pro 2019.

**Data and code availability**

All data and codes used for the electrical circuit model are available upon a reasonable request from the corresponding author (T.P.M.).

#### Supplementary Figures

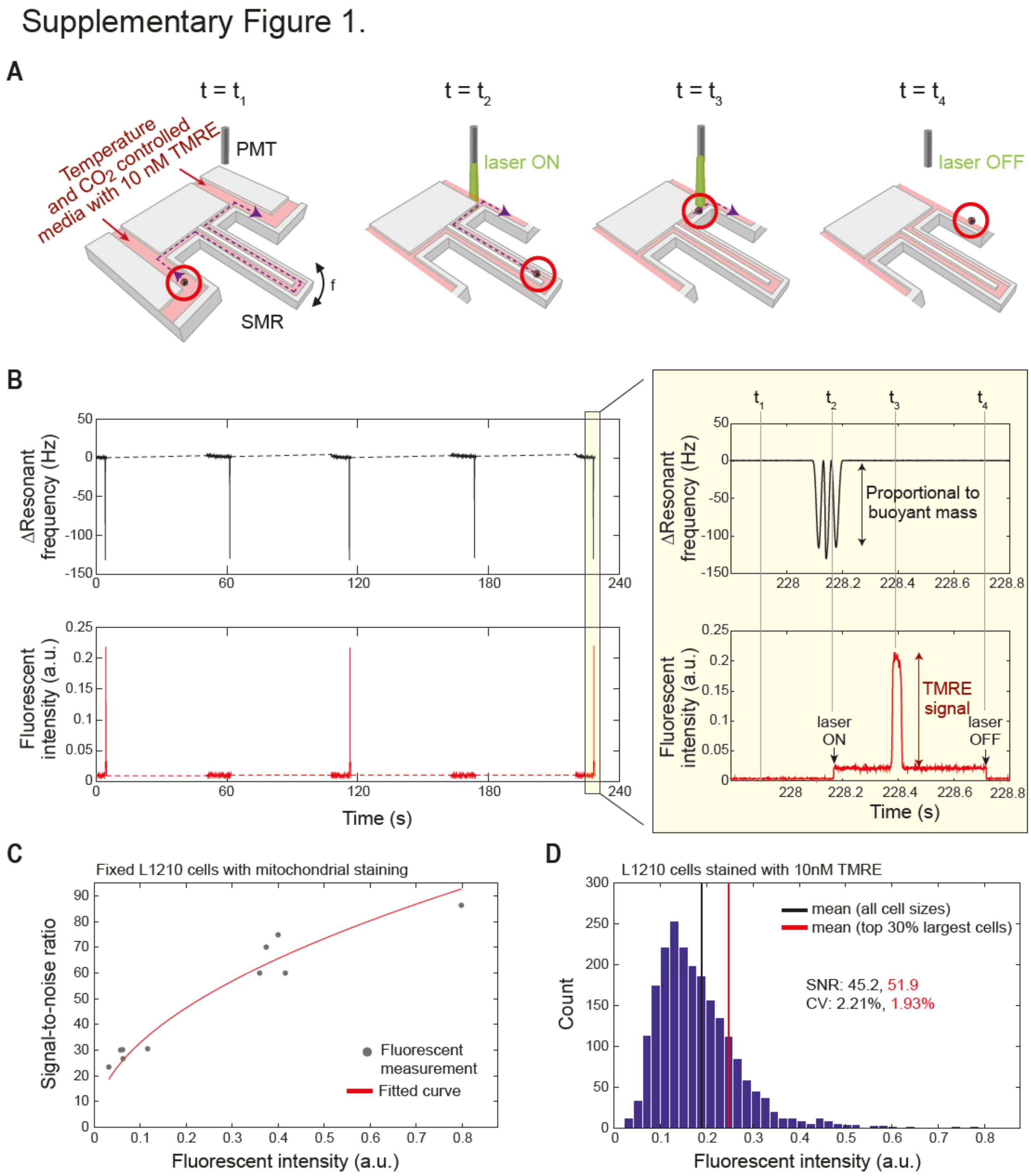

**Supplementary Fig. 1: Fluorescence detection system coupled to suspended microchannel resonator (SMR) allows precise mass-normalized TMRE signal quantifications**

**A)** Schematic of the SMR measurement setup. A cell (highlighted by red circles) is grown in environmentally controlled conditions in normal cell culture media within the SMR [1-3]. Approximately every 1 min, the cell is flown through a microchannel inside the SMR cantilever (t

= $t_1$, the trajectory of the cell is shown with purple dotted lines). This results in a change in the cantilever's resonance frequency that is proportional to the buoyant mass of the cell (t = $t_2$). The change in resonance frequency turns on a laser that is focused to the microchannel outside the SMR cantilever. As the cell passes through the laser excitation (t = $t_3$), the emission signal from the cell is captured using photomultiplier tubes (PMTs). Then, the laser turns off and the fluid flow is stopped for approximately 1 minute, after which the cell is flown back to the other side.

**B)** Example raw data of SMR resonant frequency and PMT based fluorescent detection during five consecutive buoyant mass measurements (*left*) along with a zoom-in of a single measurement of buoyant mass and TMRE signal (*right*). When a cell goes through the SMR its buoyant mass (double black arrow; height of side resonant frequency peak) acts as a trigger to turn on the laser. Approximately 200 ms later the cell flows through the laser illumination where the TMRE signal (red double arrow; height of the PMT voltage spike) is quantified using the PMTs. Approximately 200 ms later the laser turns off to minimize phototoxicity from scattered light. In data analysis each TMRE signal intensity is normalized to the corresponding buoyant mass measurement to account for cell size-dependent differences.

**C)** Quantification of fluorescent error for mitochondrial staining. Mitochondria in L1210 cells were stained with 100 nM or 333 nM MitoTracker Red CMXRos, the cells were fixed and then each cell was repeatedly measured to quantify fluorescence errors. Signal-to-noise ratio is displayed for each cell (gray dots, N=10 cells, each repeatedly measured >50 times). Estimated signal-to-noise ratio (SNR) assuming perfect shot noise (i.e., SNR $\propto \sqrt{n}$, where n is the average number of photons detected on PMT per unit time) is displayed in red.

**D)** Estimated SNR and Coefficient of Variation (CV) of typical TMRE measurement. Fluorescent Intensity of L1210 cells treated with 10nM TMRE and their distribution is shown in the blue histogram. Estimated SNR and CV of the TMRE measurements is shown for average cells in a population (black) and for top 30% largest cells in a population, which reflect mostly G2 and mitotic cells (red). SNR and CV were calculated using the fitted curve displayed in panel C.

#### Supplementary Figure 2.

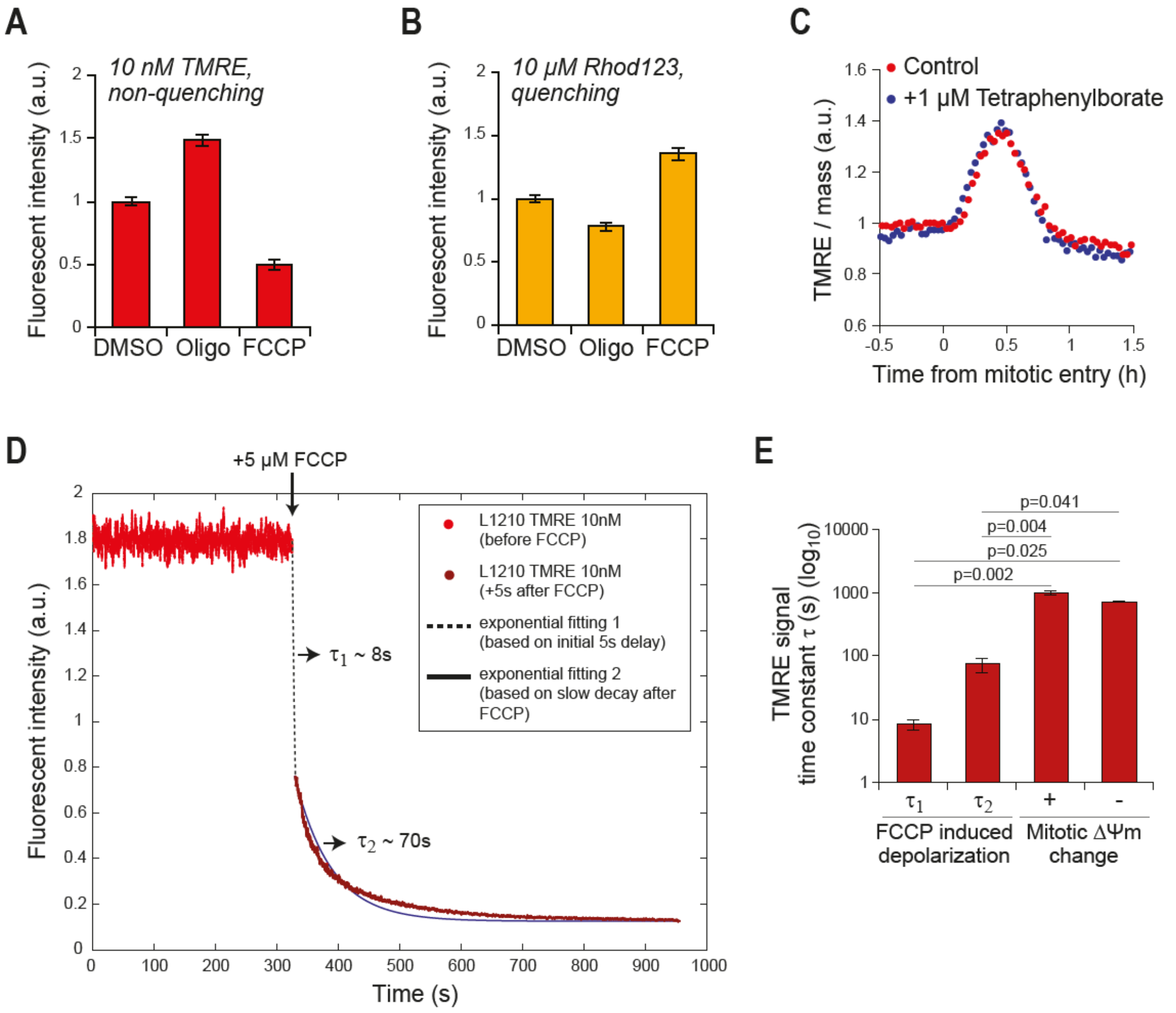

**Supplementary Fig. 2: Detection of the mitochondrial hyperpolarization at the end of the cell cycle is not limited by TMRE diffusion kinetics when TMRE is used in non-quenching mode**

**A)** Population average TMRE fluorescence intensity for L1210 cells after 30 min treatment 1 μM oligomycin or 1 μM FCCP. The directionality of the fluorescence change indicates that TMRE is acting in a non-quenching mode. Comparable results were seen with 5 nM TMRE. (N=4).

**B)** Population average Rhod123 fluorescence intensity for L1210 cells after 30 min treatment 1 μM oligomycin or 1 μM FCCP. The directionality of the fluorescence change indicates that Rhod123 is acting in a quenching mode. Note that Rhod123 was removed from the culture media prior to the addition of the chemical. (N=3).

**C)** Mass-normalized TMRE (10 nM) traces for a L1210 cell around cell division in the presence and absence of 1 μM Tetraphenylborate, which promotes the diffusion of TMRE through membranes [7,8].

**D)** Example data used for quantifications of TMRE diffusion kinetics (N=3 independent experiments). L1210 cells were stained with 10 nM TMRE and the average population TMRE intensity was sampled using flow cytometer before (red) and after (orange) treatment with the protonophore FCCP (5 μM). The time constants were extracted by i) fitting an exponential decrease between the last data point prior to FCCP addition and the first data point after FCCP addition ($\tau_1$), or by ii) fitting an exponential decrease to the data after FCCP addition ($\tau_2$).

**E)** Quantifications of TMRE signal time constants during FCCP induced depolarization ($\tau_1$ and $\tau_2$ from panel D, N=3) and during the TMRE increase (+) and decrease (-) in mitosis (N=40, n=85). Note that the rate at which TMRE signal changes in mitosis is at least an order of magnitude slower than what TMRE signal can change. p-values were obtained using ANOVA followed by Sidakholm test.

#### Supplementary Figure 3.

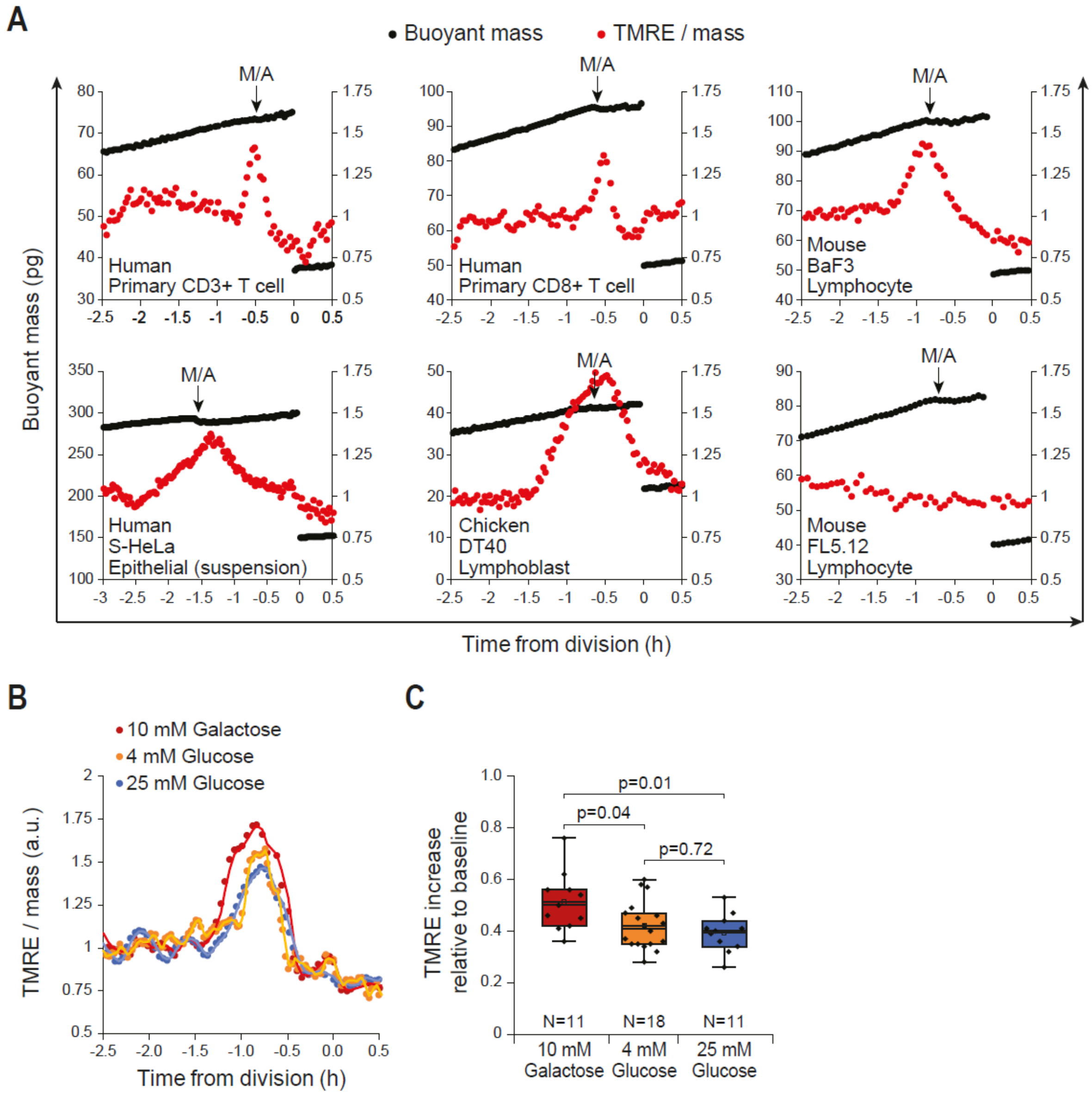

**Supplementary Fig. 3: Mitochondrial hyperpolarization takes place at the end of cell cycle in various animal cell types and under different glucose concentrations.**

**A)** Buoyant mass (black) and mass-normalized TMRE (red) traces for indicated primary cells and cell lines. Metaphase-anaphase transition (M/A) is indicated for each cell with black arrow. Note that while primary cells and several cell lines display mitochondrial hyperpolarization at the end of cell cycle, FL5.12 lymphocytes do not, indicating that mitochondrial hyperpolarization is not a general requirement for cell division.

**B)** Mass-normalized TMRE traces for L1210 cells around cell division under indicated glucose conditions. Galactose grown cells did not have glucose in the culture media.

**C)** Quantifications of L1210 TMRE increase in indicated glucose conditions. p-values were obtained using ANOVA followed by Sidakholm test.

Supplementary Figure 4.

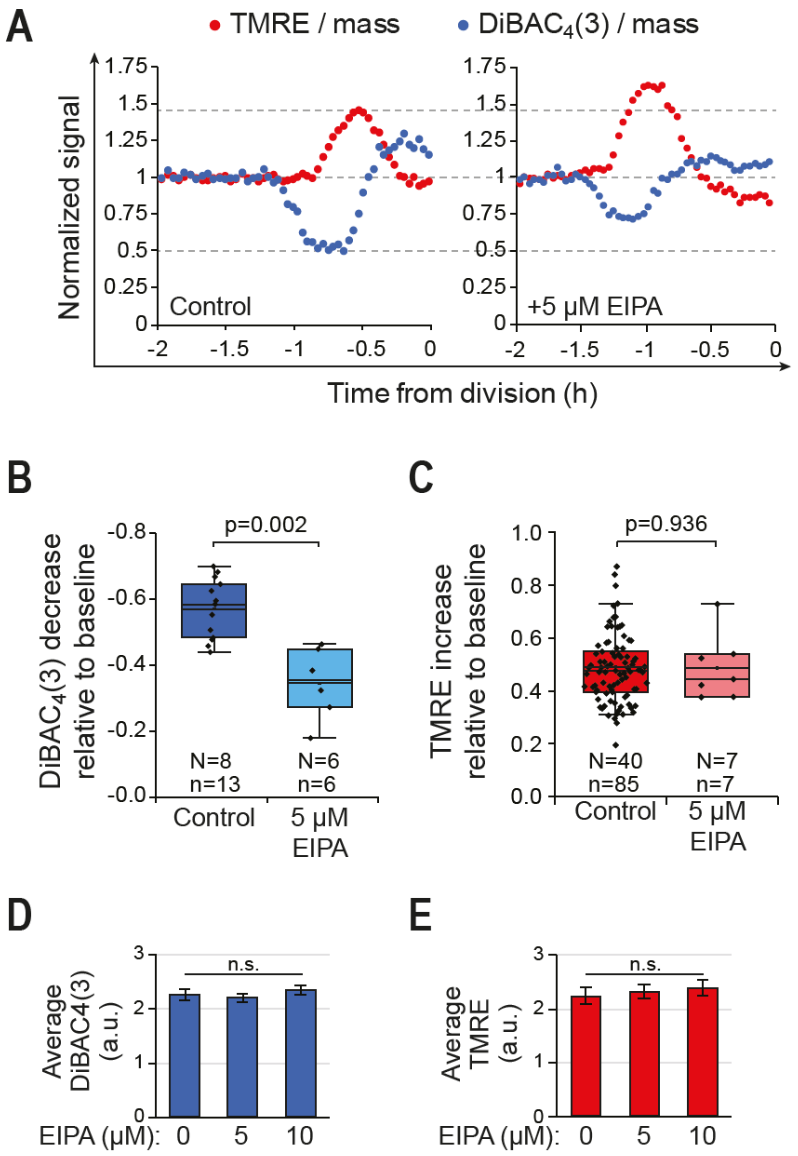

**Supplementary Fig. 4: Plasma membrane hyperpolarization is coupled to mitotic cell swelling with little influence on TMRE signal.**

**A)** Mass-normalized TMRE (red, 10 nM) and DiBAC$_4$(3) (blue, 1µM) traces for a control (left) and 5 µM EIPA treated (right) L1210 cell around cell division. Note that the DiBAC$_4$(3) signal decrease and TMRE signal increase are not starting or ending at the same time. Also, the EIPA treatment, which inhibits mitotic cell swelling [1], reduces the DiBAC$_4$(3) signal change but not the TMRE signal change, indicating that the plasma membrane potential changes are not significantly affecting the TMRE signal.

**B)** Quantification of DiBAC$_4$(3) changes shown in panel A. p-value obtained using two-tailed Welch's t-test.

**C)** Quantification of TMRE changes shown in panel A. p-value obtained using two-tailed Welch's t-test.

**D)** Quantification of population average DiBAC$_4$(3) signal after a treatment with EIPA. p-value obtained using ANOVA (N=3).

**E)** Quantification of population average TMRE signal after a treatment with EIPA. p-value obtained using ANOVA (N=3).

#### Supplementary Figure 5.

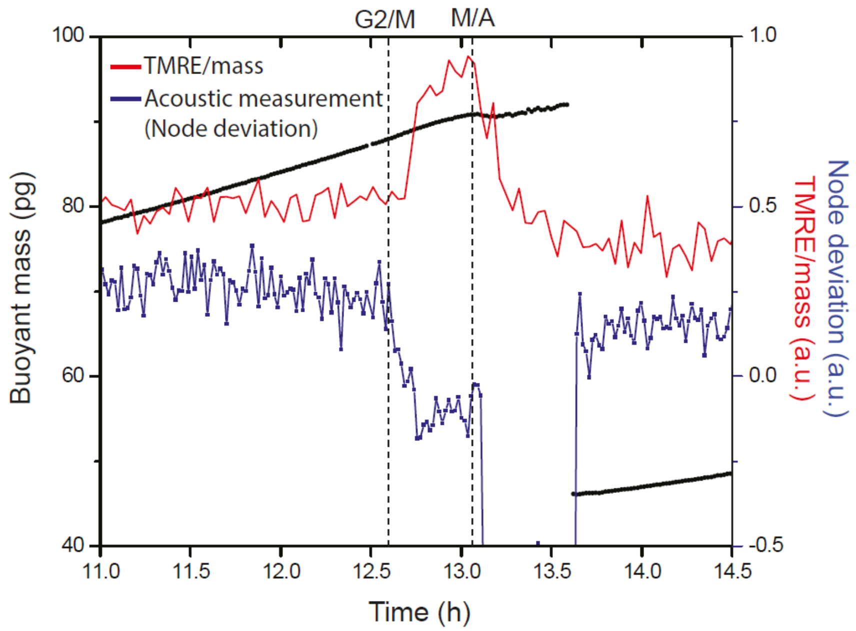

**Supplementary Fig. 5: TMRE signal increases after G2/M transition until the metaphase-anaphase transition.**

Buoyant mass (black), mass-normalized TMRE (red) and node deviation (acoustic measurement dependent on cell stiffness, blue) trace for a L1210 cell around cell division. The node deviation decreases slightly after mitotic entry as cells swell up and node deviation rapidly drops as cells elongate in anaphase, thus allowing detection of G2/M transition and metaphase-anaphase transition [5].

### Supplementary Figure 6.

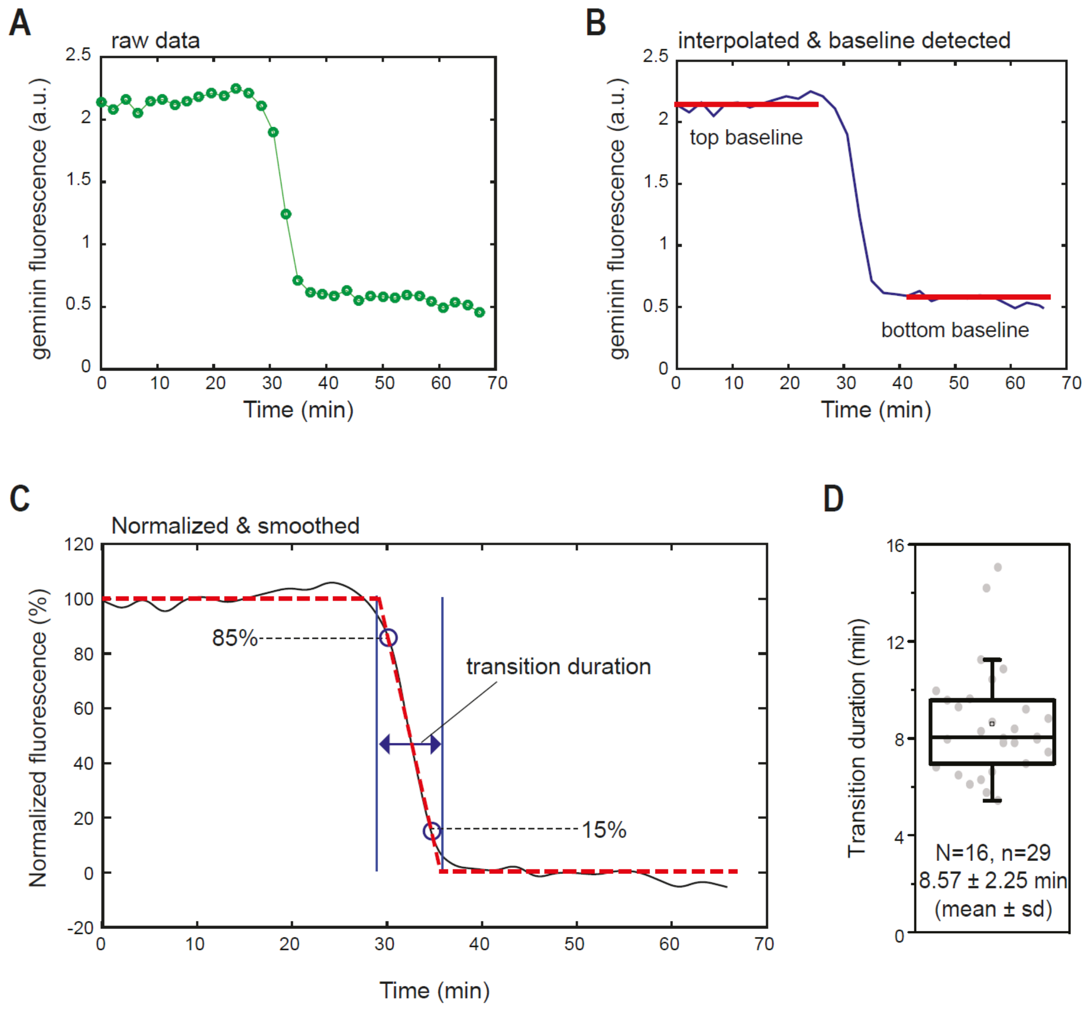

**Supplementary Fig. 6: Protein degradation at metaphase-anaphase transition takes approximately 8 min in L1210 cells.**

**A)** mAG-Geminin fluorescence trace for a L1210 FUCCI cell around metaphase-anaphase transition. mAG-Geminin is degraded at metaphase-anaphase transition due to APC/C activity.

**B)** Data in panel A after data interpolation. Averaged mAG-Geminin fluorescence level before (top baseline) and after (bottom baseline) metaphase-anaphase transition is also indicated in red.

**C)** Data in panel B after normalization and data smoothing. This data was used to detect the transition duration by fitting a straight line between 85% and 15% mAG-Geminin fluorescence intensity points (dark blue circles) and detecting the time points where this straight line crosses the top and bottom baselines.

**D)** Duration of protein degradation at the metaphase-anaphase transition quantified based mAG-Geminin degradation in 29 L1210 FUCCI cells.

#### Supplementary Figure 7.

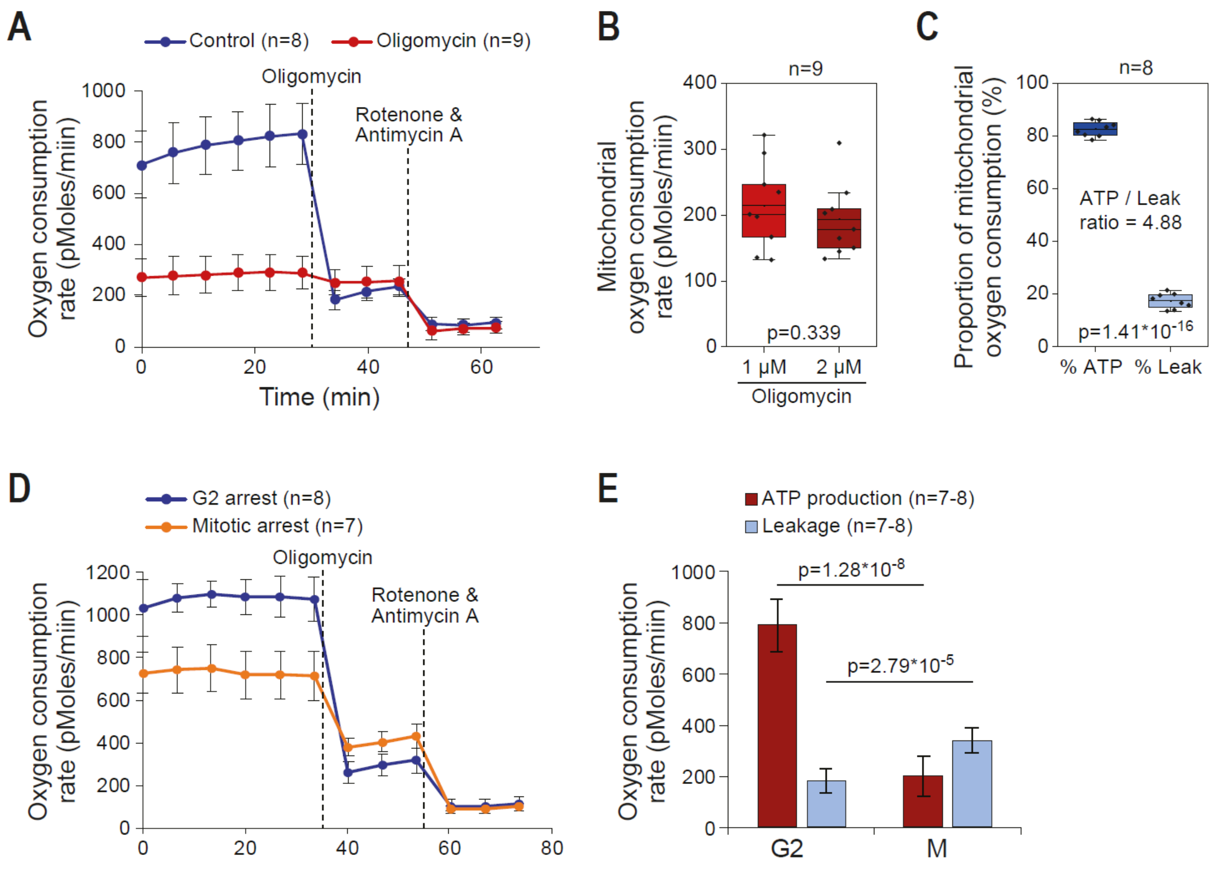

**Supplementary Fig. 7: L1210 cells arrested in mitosis display lower mitochondrial ATP synthesis rates than cells arrested in G2.**

**A)** Oxygen consumption rate measurements of control L1210 cells and in L1210 cells pre-treated with 1 µM oligomycin for 4 h. Both samples were injected with 1 µM oligomycin and later with 2 µM Rotenone & 2 µM Antimycin A, as indicated by dashed lines.

**B)** Quantification of oxygen consumption rates in the 1 µM oligomycin pretreated L1210 cells before and after injection of additional oligomycin, as shown in panel A. Note that the oxygen consumption doesn't decrease further by additional oligomycin, indicating that 1 µM oligomycin is adequate to achieve maximal inhibition of ATP synthase. p-value obtained using two-tailed Welch's t-test.

**C)** Quantification of proportional oxygen consumption rates attributable to leakage and ATP synthesis in control L1210 cells, as shown in panel A. p-value obtained using two-tailed Welch's t-test.

**D)** Oxygen consumption rate measurements of L1210 cells arrested to G2 (blue) or to prometaphase (mitotic arrest, orange). Both samples were injected with 1 µM oligomycin and later with 2 µM Rotenone & 2 µM Antimycin A, as indicated by dashed lines. The mitotic arrest was obtained by releasing G2 arrested cells to media with STLC (kinesin motor inhibitor that arrests

the cells to prometaphase) 2h prior to measurement. Note that mitotic arrests inhibit cell growth [3] and result in mitochondrial degradation [40], which may affect the oxygen consumption rates.
**E)** Quantification of ATP synthesis and leakage rates in the G2 and mitotic arrested L1210 cells, based on the oxygen consumption measurements shown in panel D. p-values obtained using two-tailed Welch's t-test.

### Supplementary Figure 8.

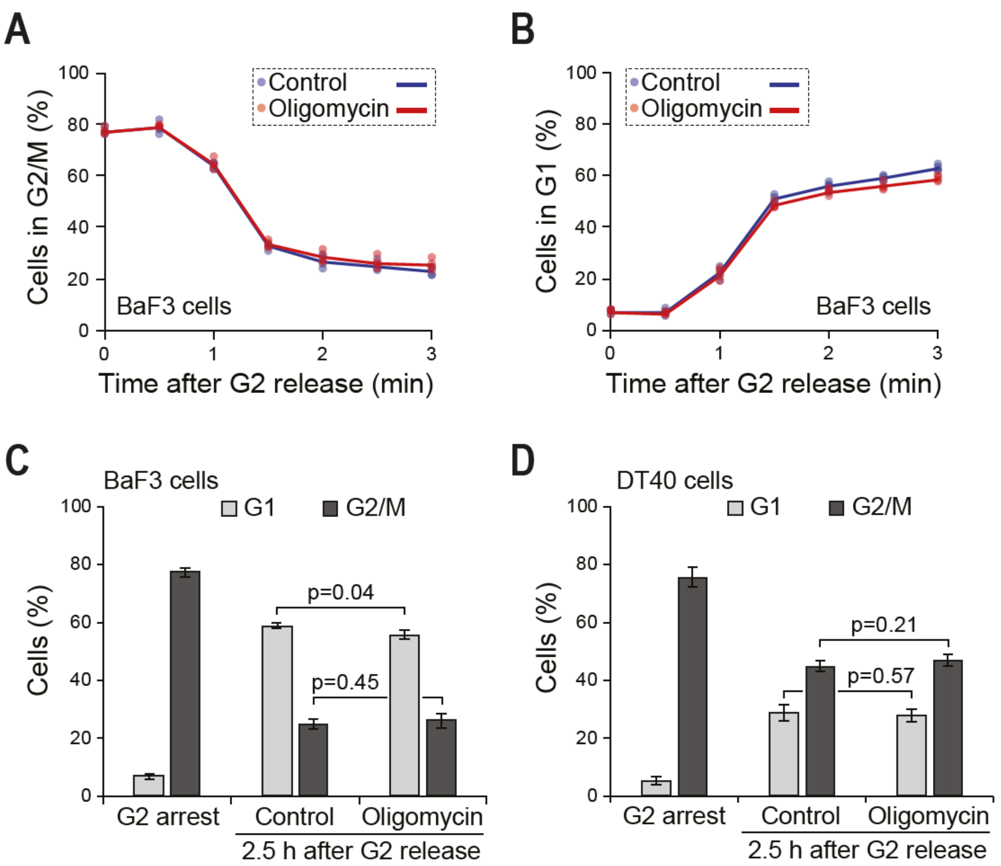

**Supplementary Fig. 8: Inhibition of mitochondrial ATP synthesis has little effect on mitotic progression.**

**A)** Relative number of G2/M cells in control (blue) and 1 µM oligomycin (red) treated BaF3 cells at indicated times after release from G2 arrest. Oligomycin treatment was started 15 min before release from G2 and was maintained on cells after G2 release. Each dot represents a separate culture (n=4).

**B)** Same as panel A, except data is indicating the relative number of G1 cells at indicated times after release from G2 arrest.

**C)** Quantifications of BaF3 data in panels A and B for the 2.5 h timepoint after G2 release (n=4 separate cultures). p-values obtained using two-tailed Welch's t-test.

**D)** Same as panel C, except experiments were carried out with DT40 chicken lymphocytes (n=4 separate cultures). p-values obtained using two-tailed Welch's t-test.

### Supplementary Figure 9.

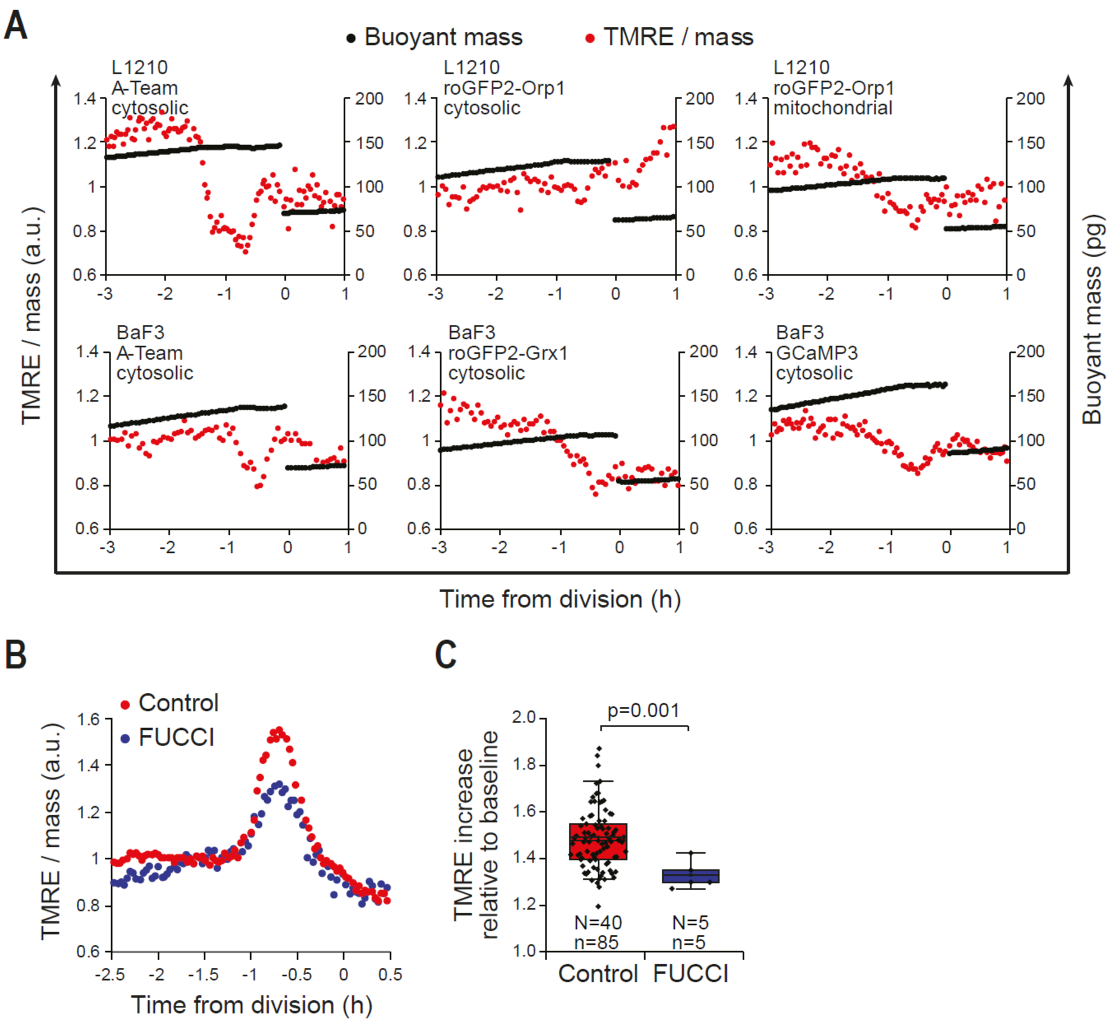

**Supplementary Fig. 9: Mitotic mitochondrial hyperpolarization is perturbed by expression of exogenous proteins.**

**A)** Buoyant mass (black) and mass-normalized TMRE (red) traces for L1210 and BaF3 cells around cell division. The cells were constitutively expressing indicated genetic constructs (FRET based fluorescent sensors for various metabolites) and the TMRE hyperpolarization was no longer observed.

**B)** Mass-normalized TMRE traces for control (wild-type, red) and FUCCI (blue) L1210 cells around cell division.

**C)** Quantification of the mitotic TMRE increase in control (red) and FUCCI (blue) L1210 cells, as shown in panel B. Note that the FUCCI cells express mAG-Geminin under endogenous geminin promoter, and while mitochondria still hyperpolarize, this hyperpolarization is less extensive than in control (wild-type) L1210 cells. p-value obtained using two-tailed Welch's t-test.

Supplementary Figure 10.

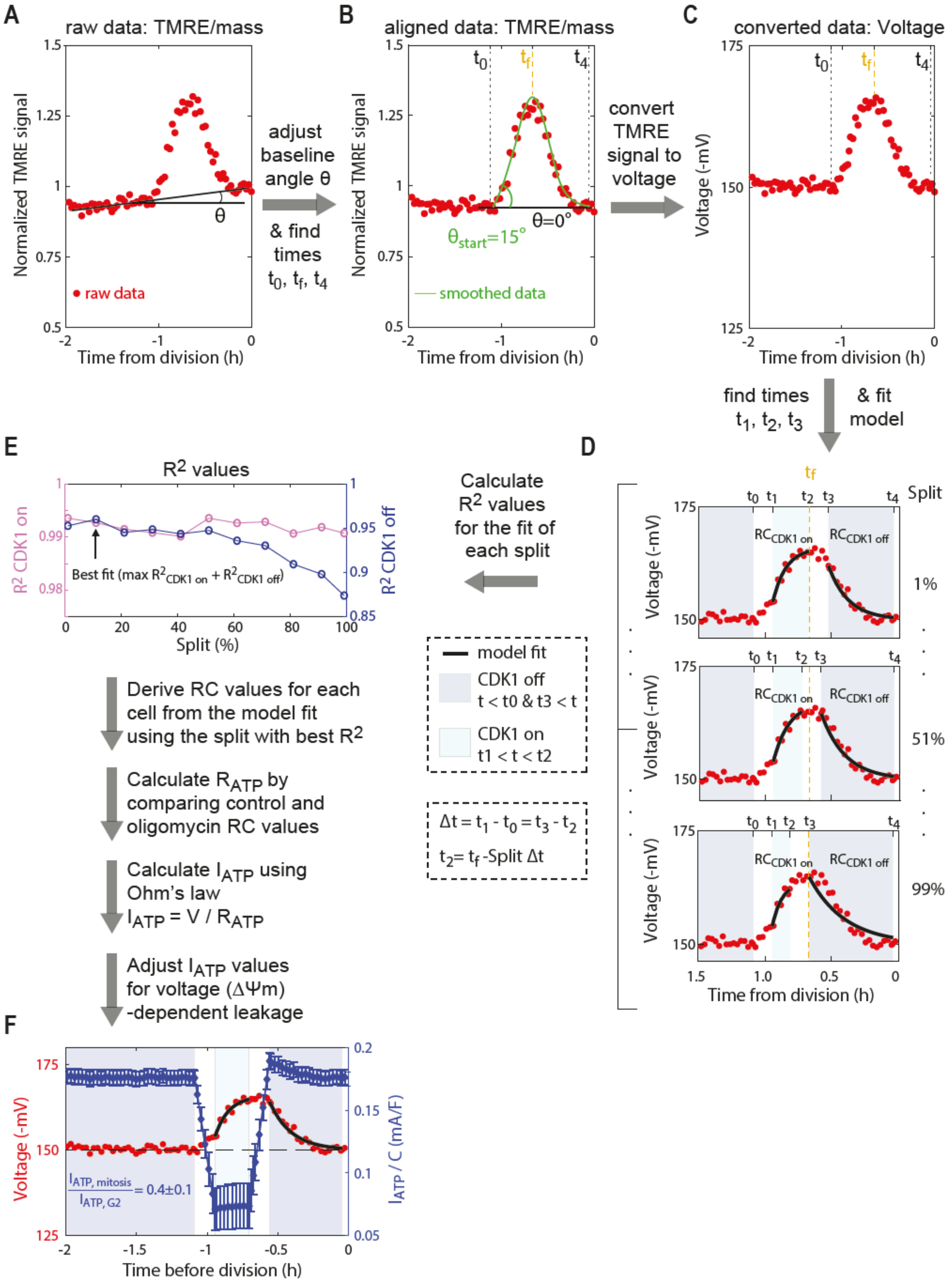

**Supplementary Fig. 10: Applying electrical circuit model to mitotic TMRE data reveals the current flowing through ATP synthase.**

Workflow for the modelling:

**A)** TMRE/mass signal around mitosis is normalized to have zero baseline slope (θ). Note that the baseline slope varied between cells and slope corrections were done to both directions.

**B)** Normalized TMRE/mass signal is fitted a smoothed curve (green) with smoothing window of 20 min. Using the smoothed curve, we find the beginning of the hyperpolarization at time $t_0$ where start slope is $\theta_{start} = 15^o$, the maximum of TMRE at time $t_f$ and the end of depolarization at time $t_4$ where TMRE has returned to baseline (G2) levels. Cells that divided before a new baseline was clearly established were excluded from the modelling.

**C)** TMRE/mass signal is converted to approximate voltage according to previously reported scaling between tetramethylrhodamidine ester fluorescence signal and membrane potentials [11]. $V(mV) = (TMRE/c_1)/c_2$, where $c_1 = 29.109$ and $c_2 = 0.0215$. Baseline voltage was set to -150 mV.

**D)** Voltage signal is divided into four periods where: i) CDK1 is off (dark blue, $t_0 < t \ \& \ t > t_3$), ii) CDK1 transitions from off to on (white, $t_0 < t < t_1$ ), iii) CDK1 is on (light blue, $t_1 < t < t_2$ ), and iv) CDK1 transitions from off to on (white, $t_2 < t < t_3$ ). The transitions are assumed to have fixed durations ($\Delta t = t_3 - t_2 = t_1 - t_0 = 8.57\ min$), obtained by quantifying the duration of geminin protein degradation at metaphase-anaphase transition (Figure S6). Since $t_2$ (marking end of period CDK1 where is on) is expected to be near the approximate metaphase-anaphase transition ($t_f$), yet the precise location of $t_2$ is unknown, eleven values $Split = 1,11,21, \dots ,91,99\ \%$ were used to obtain eleven corresponding values $t_2 = t_f - Split\ \Delta t$. For each one of these values, the CDK1 on and CDK1 off periods were fitted with analytical solutions to the electrical circuit model (Figure 4A) in order to obtain corresponding $RC$ values.

**E)** The quality of the model fitting is estimated by calculating $R^2_{CDK1\ on}$ and $R^2_{CDK1\ off}$ values for each Split. For further analysis, the RC values of each cell are derived from the Split with best fit overall for the specific cell. This procedure is repeated for each control and oligomycin treated cell to obtain population averages for $RC_{CDK1\ on}$ and $RC_{CDK1\ off}$ values, which are then used to back calculate $R_{ATP}$ for CDK1 on and off states. The $R_{ATP}$ values are then applied to each voltage converted TMRE trace to quantify the current flowing through ATP synthase, i.e. ATP synthesis rate.

**F)** Example of voltage (ΔΨm, red) and ATP synthesis rate ($I_{ATP}/C$, blue) for a single L1210 cells around cell division. Error bars depict the s.e.m. of the $R_{ATP}$.

#### Supplementary Figure 11.

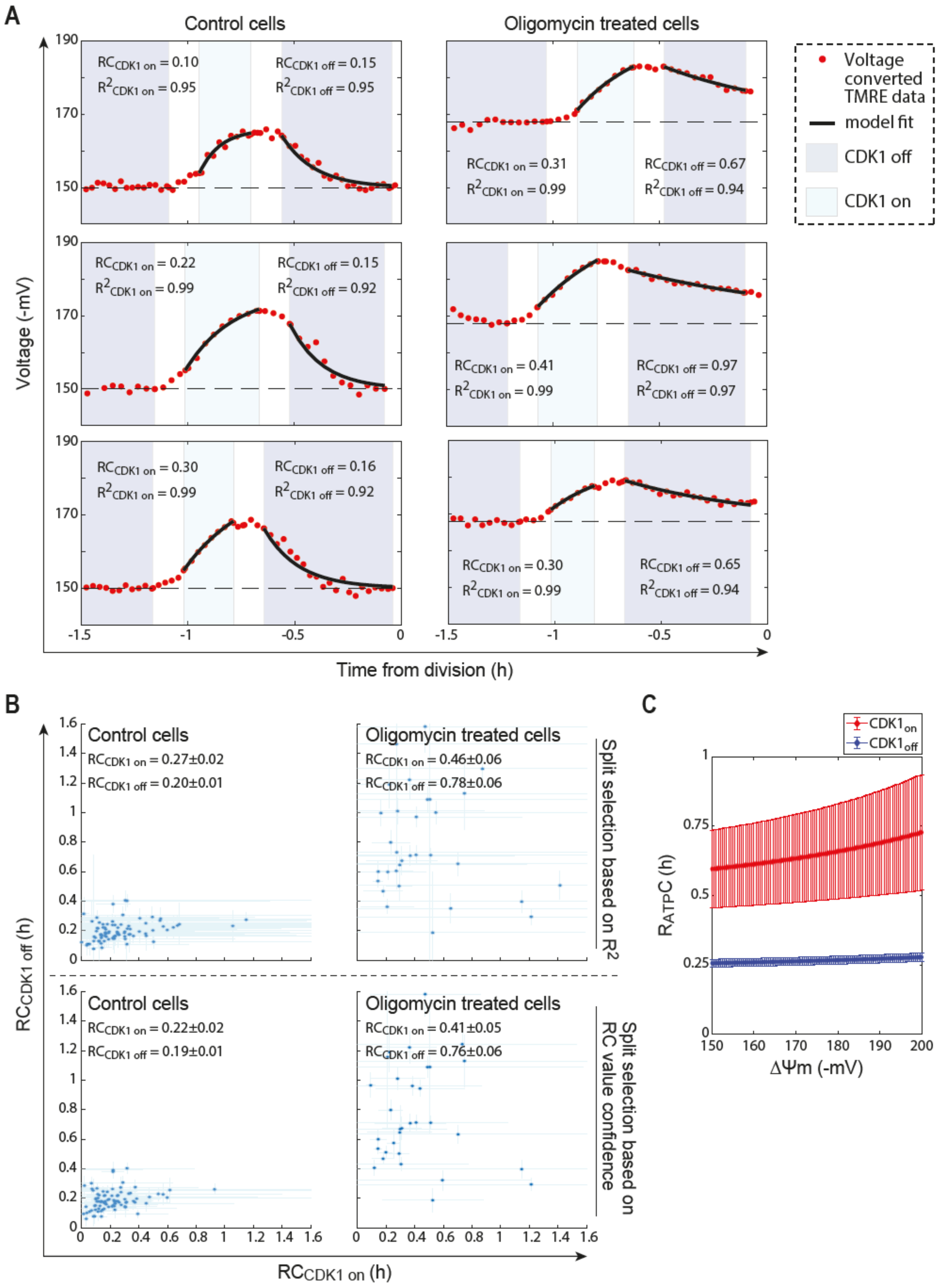

**Supplementary Fig. 11: Oligomycin treatment results in increased RC values during late mitosis**

**A)** Control (*left*) and 1 µM oligomycin (*right*) treated L1210 cell voltage traces. RC and $R^2$ values derived by fitting the model analytical solution (black line) to the CDK1 on (light blue) and CDK1 off (dark blue) regions are indicated.

**B)** Summary of the RC values derived for each control and oligomycin treated cell when the fitting regions (Split) were selected by maximizing $R^2$ (*top*) or by minimizing confidence interval of the derived RC (*bottom*). Each blue dot represents a single cell and error bars depict the confidence interval of the derived RC value. The written RC values depict population mean ± s.e.m.

**C)** The $R_{ATP}$ values derived for *CDK1 on* (red) and *CDK1 off* (blue) states as a function of ΔΨm. The calculated $R_{ATP}$ values depend on ΔΨm due to the non-ohmic leakage scaling with ΔΨm. Data depicts mean ± s.e.m. (n=85 for control cells, 32 for oligomycin treated cells).

#### Supplementary Figure 12.

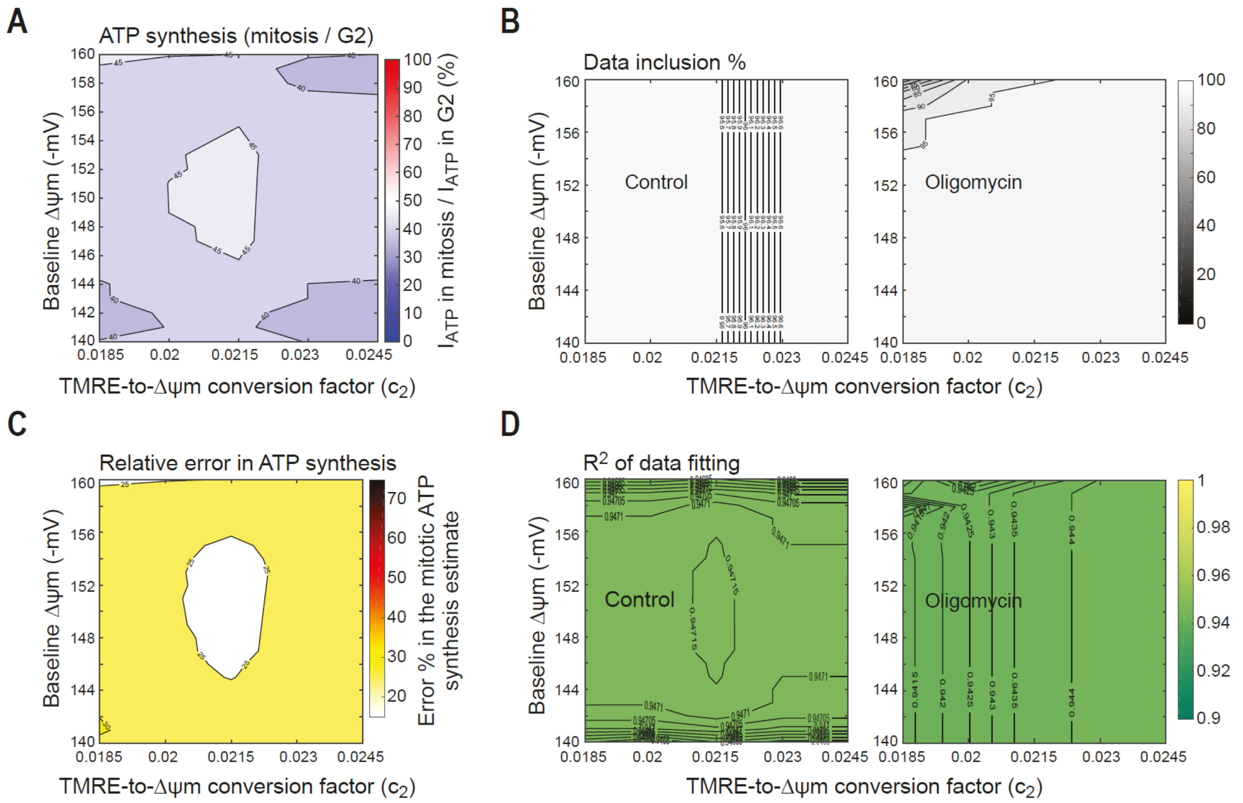

**Supplementary Fig. 12: Modeling of mitotic ATP synthesis rates is not sensitive to the values used for TMRE-to-voltage conversion.**

Sensitivity analysis over parameters of TMRE-to-ΔΨm conversion factor ($c_2 = 0.0185, 0.02, 0.0215, 0.0230, 0.0245$) and baseline voltage ($\Delta\Psi m = 140, 151, \ldots, 160$). The parameters for estimating the ΔΨm-dependent leakage were held fixed ($\beta = 4.88, \gamma = 0.0122$), as were the parameters used to define fitting regions, the fitting start angle and the duration of CDK1 on/off transition ($\theta_{start} = 15\ degrees, \Delta t = 8.57 min$).

**A)** Contour plot of $r_{ATP} = I_{ATP\ in\ mitosis}/I_{ATP\ in\ G2}$. Sensitivity analysis around reference values ($c_2 = 0.0215, \Delta\Psi m = 150\text{mV}$) showed $r_{ATP} = 46\%$, whereas varying $c_2$ or baseline voltage resulted in $r_{ATP}$ range of $35 - 50\%$.

**B)** Data inclusion % for control (left) and oligomycin (right) datasets.

**C)** Relative error $\delta(I_{ATP\ in\ mitosis}/I_{ATP\ in\ G2})\ /\ (I_{ATP\ in\ mitosis}/I_{ATP\ in\ G2})$. Sensitivity analysis around reference values ($c_2 = 0.0215, \Delta\Psi m = 150\text{mV}$) showed relative error $< 25\%$, whereas varying $c_2$ or baseline voltage resulted in errors increasing up to 30%.

**D)** Fitting quality ($R^2$) for control (left) and oligomycin (right) datasets.

#### Supplementary Figure 13.

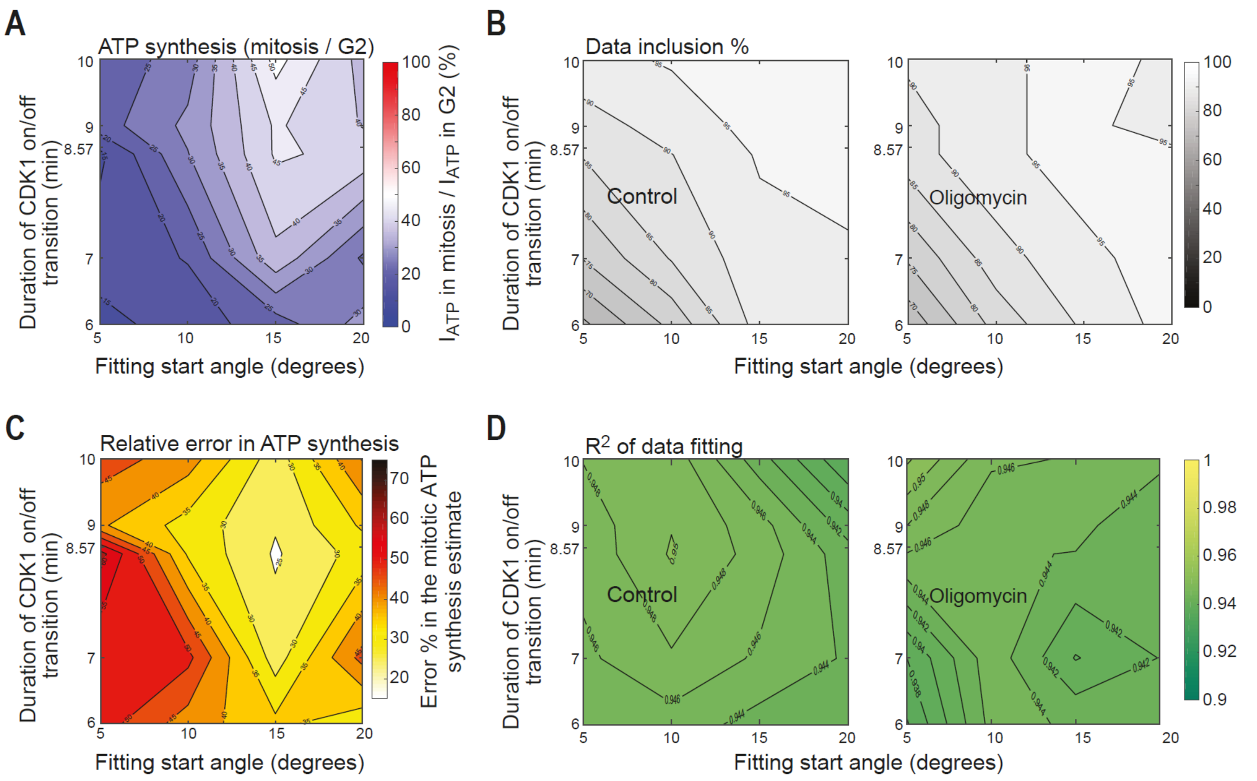

**Supplementary Fig. 13: Changing fitting regions used to derive RC values results in a model prediction with more pronounced inhibition of ATP synthesis in mitosis**

Sensitivity analysis over parameters used to define fitting regions, the fitting start angle ($\theta_{start} = 5, 10, 15, 20\ degrees$) and duration of CDK1 on/off transition ($\Delta t = 6, 7, 8.57, 9, 10\ min$). The parameters for the TMRE-to-ΔΨm conversion factor and baseline voltage were held fixed ($c_2 = 0.0215, \Delta\Psi m = 150$ mV), as were the parameters for estimating the ΔΨm-dependent leakage ($\beta = 4.88, \gamma = 0.0122$).

**A)** Contour plot of $r_{ATP} = I_{ATP\ in\ mitosis}/I_{ATP\ in\ G2}$. Sensitivity analysis around reference values ($\theta_{start} = 15\ degrees$ and $\Delta t = 8.57\ min$) showed $r_{ATP} = 46\%$, whereas varying $\theta_{start}$ or $\Delta t$ resulted in $r_{ATP}$ range of $15 - 50\%$.

**B)** Data inclusion % for control (left) and oligomycin (right) datasets.

**C)** Relative error $\delta(I_{ATP\ in\ mitosis}/I_{ATP\ in\ G2})\ /\ (I_{ATP\ in\ mitosis}/I_{ATP\ in\ G2})$. Sensitivity analysis around reference values ($c_2 = 0.0215, \Delta\Psi m = 150$mV) showed relative error $< 25\%$, whereas varying $\theta_{start}$ or $\Delta t$ resulted in errors increasing up to 55%.

**D)** Fitting quality ($R^2$) for control (left) and oligomycin (right) datasets.

### Supplementary Figure 14.

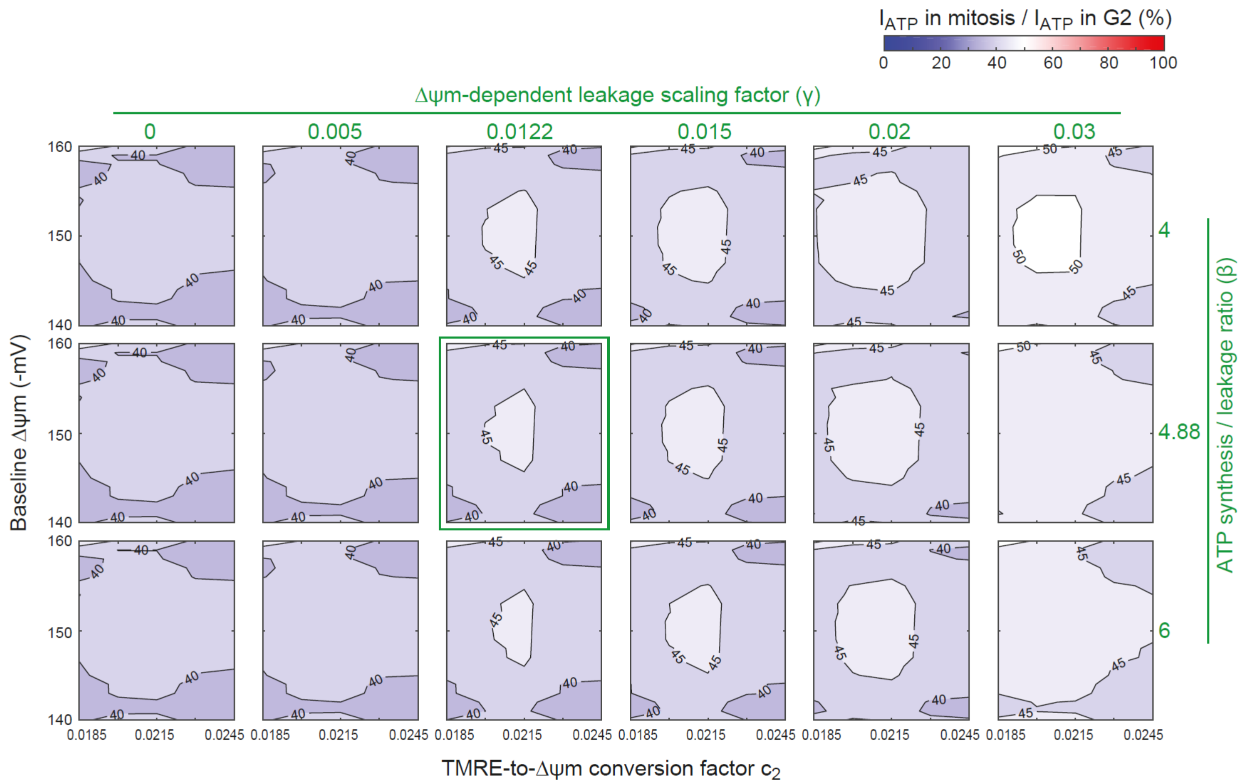

**Supplementary Fig. 14: Modeling of mitotic ATP synthesis rates has little sensitivity to the values used for estimating the non-ohmic scaling of ΔΨm-dependent leakage.**
Sensitivity analysis over parameters used for estimating the non-ohmic scaling of ΔΨm-dependent leakage ($\beta = 4, 4.88, 6$ *and* $\gamma = 0, 0.005, 0.0122, 0.015, 0.02, 0.03$). The parameters used to define fitting regions, the fitting start angle and the duration of CDK1 on/off transition ($\theta_{start} = 15\ degrees, \Delta t = 8.57min$), were held fixed. The parameters for the TMRE-to-ΔΨm conversion were varied in similar range as in Figure S12. Data depicts contour plots of $r_{ATP} = I_{ATP\ in\ mitosis}/I_{ATP\ in\ G2}$. The green box indicates the reference values ($\beta = 4.88, \gamma = 0.0122$). Overall, varying parameters used for estimating the non-ohmic scaling of ΔΨm-dependent leakage resulted in $r_{ATP}$ range of $35 - 55\%$.

#### Supplementary References

1 Son, S. *et al.* Resonant microchannel volume and mass measurements show that suspended cells swell during mitosis. *J Cell Biol* **211**, 757-763, doi:10.1083/jcb.201505058 (2015).

2 Son, S. *et al.* Direct observation of mammalian cell growth and size regulation. *Nat Methods* **9**, 910-912, doi:10.1038/nmeth.2133 (2012).

3 Miettinen, T. P., Kang, J. H., Yang, L. F. & Manalis, S. R. Mammalian cell growth dynamics in mitosis. *Elife* **8**, doi:10.7554/eLife.44700 (2019).

4 Burg, T. P. *et al.* Weighing of biomolecules, single cells and single nanoparticles in fluid. *Nature* **446**, 1066-1069, doi:10.1038/nature05741 (2007).

5 Kang, J. H. *et al.* Noninvasive monitoring of single-cell mechanics by acoustic scattering. *Nat Methods* **16**, 263-269, doi:10.1038/s41592-019-0326-x (2019).

6 Perry, S. W., Norman, J. P., Barbieri, J., Brown, E. B. & Gelbard, H. A. Mitochondrial membrane potential probes and the proton gradient: a practical usage guide. *Biotechniques* **50**, 98-115, doi:10.2144/000113610 (2011).

7 Farkas, D. L., Wei, M. D., Febbroriello, P., Carson, J. H. & Loew, L. M. Simultaneous imaging of cell and mitochondrial membrane potentials. *Biophys J* **56**, 1053-1069, doi:10.1016/S0006-3495(89)82754-7 (1989).

8 Ward, M. W., Rego, A. C., Frenguelli, B. G. & Nicholls, D. G. Mitochondrial membrane potential and glutamate excitotoxicity in cultured cerebellar granule cells. *J Neurosci* **20**, 7208-7219 (2000).

9 Nicholls, D. G. & Ward, M. W. Mitochondrial membrane potential and neuronal glutamate excitotoxicity: mortality and millivolts. *Trends Neurosci* **23**, 166-174 (2000).

10 Ehrenberg, B., Montana, V., Wei, M. D., Wuskell, J. P. & Loew, L. M. Membrane-Potential Can Be Determined in Individual Cells from the Nernstian Distribution of Cationic Dyes. *Biophysical Journal* **53**, 785-794, doi:Doi 10.1016/S0006-3495(88)83158-8 (1988).

11 Gerencser, A. A. *et al.* Quantitative measurement of mitochondrial membrane potential in cultured cells: calcium-induced de- and hyperpolarization of neuronal mitochondria. *J Physiol* **590**, 2845-2871, doi:10.1113/jphysiol.2012.228387 (2012).

12 Zorova, L. D. *et al.* Functional Significance of the Mitochondrial Membrane Potential. *Biochem Mosc Suppl S* **12**, 20-26, doi:10.1134/S1990747818010129 (2018).

13 Zlotek-Zlotkiewicz, E., Monnier, S., Cappello, G., Le Berre, M. & Piel, M. Optical volume and mass measurements show that mammalian cells swell during mitosis. *J Cell Biol* **211**, 765-774, doi:10.1083/jcb.201505056 (2015).

14 Miettinen, T. P. *et al.* Identification of transcriptional and metabolic programs related to mammalian cell size. *Curr Biol* **24**, 598-608, doi:10.1016/j.cub.2014.01.071 (2014).

15 Miettinen, Teemu P. & Björklund, M. Cellular Allometry of Mitochondrial Functionality Establishes the Optimal Cell Size. *Developmental Cell* **39**, doi:10.1016/j.devcel.2016.09.004 (2016).

16 Kitami, T. *et al.* A Chemical Screen Probing the Relationship between Mitochondrial Content and Cell Size. *Plos One* **7**, doi:ARTN e33755 10.1371/journal.pone.0033755 (2012).

17 Gan, Z., Audi, S. H., Bongard, R. D., Gauthier, K. M. & Merker, M. P. Quantifying mitochondrial and plasma membrane potentials in intact pulmonary arterial endothelial cells based on extracellular disposition of rhodamine dyes. *Am J Physiol-Lung C* **300**, L762-L772, doi:10.1152/ajplung.00334.2010 (2011).

18 Nicholls, D. G. & Budd, S. L. Mitochondria and neuronal survival. *Physiol Rev* **80**, 315-360, doi:10.1152/physrev.2000.80.1.315 (2000).

19 Gutscher, M. *et al.* Proximity-based protein thiol oxidation by H2O2-scavenging peroxidases. *J Biol Chem* **284**, 31532-31540, doi:10.1074/jbc.M109.059246 (2009).

20 Gutscher, M. *et al.* Real-time imaging of the intracellular glutathione redox potential. *Nat Methods* **5**, 553-559, doi:10.1038/nmeth.1212 (2008).

21 Kotera, I., Iwasaki, T., Imamura, H., Noji, H. & Nagai, T. Reversible dimerization of Aequorea victoria fluorescent proteins increases the dynamic range of FRET-based indicators. *ACS Chem Biol* **5**, 215-222, doi:10.1021/cb900263z (2010).

22 Imamura, H. *et al.* Visualization of ATP levels inside single living cells with fluorescence resonance energy transfer-based genetically encoded indicators. *Proc Natl Acad Sci U S A* **106**, 15651-15656, doi:10.1073/pnas.0904764106 (2009).

23 Tian, L. *et al.* Imaging neural activity in worms, flies and mice with improved GCaMP calcium indicators. *Nat Methods* **6**, 875-881, doi:10.1038/nmeth.1398 (2009).

24 Gavet, O. & Pines, J. Progressive activation of CyclinB1-Cdk1 coordinates entry to mitosis. *Dev Cell* **18**, 533-543, doi:10.1016/j.devcel.2010.02.013 (2010).

25 Lindqvist, A., Rodriguez-Bravo, V. & Medema, R. H. The decision to enter mitosis: feedback and redundancy in the mitotic entry network. *J Cell Biol* **185**, 193-202, doi:10.1083/jcb.200812045 (2009).

26 Hegarat, N., Rata, S. & Hochegger, H. Bistability of mitotic entry and exit switches during open mitosis in mammalian cells. *Bioessays* **38**, 627-643, doi:10.1002/bies.201600057 (2016).

27 Brand, M. D. & Nicholls, D. G. Assessing mitochondrial dysfunction in cells. *Biochem J* **435**, 297-312, doi:10.1042/BJ20110162 (2011).

28 Nicholls, D. G. Mitochondrial membrane potential and aging. *Aging Cell* **3**, 35-40 (2004).

29 Dimroth, P., Kaim, G. & Matthey, U. Crucial role of the membrane potential for ATP synthesis by F1Fo ATP synthases. *J Exp Biol* **203**, 51-59 (2000).

30 Mitchell, P. & Moyle, J. Estimation of membrane potential and pH difference across the cristae membrane of rat liver mitochondria. *Eur J Biochem* **7**, 471-484, doi:10.1111/j.1432-1033.1969.tb19633.x (1969).

31 Brand, M. D. Mitochondrial generation of superoxide and hydrogen peroxide as the source of mitochondrial redox signaling. *Free Radical Bio Med* **100**, 14-31, doi:10.1016/j.freeradbiomed.2016.04.001 (2016).

32 Jastroch, M., Divakaruni, A. S., Mookerjee, S., Treberg, J. R. & Brand, M. D. Mitochondrial proton and electron leaks. *Essays Biochem* **47**, 53-67, doi:10.1042/bse0470053 (2010).

33 Nobes, C. D., Brown, G. C., Olive, P. N. & Brand, M. D. Non-ohmic proton conductance of the mitochondrial inner membrane in hepatocytes. *J Biol Chem* **265**, 12903-12909 (1990).

34 Loew, L. M., Tuft, R. A., Carrington, W. & Fay, F. S. Imaging in five dimensions: time-dependent membrane potentials in individual mitochondria. *Biophys J* **65**, 2396-2407, doi:10.1016/S0006-3495(93)81318-3 (1993).

35 Conn, A. R., Gould, N. I. M. & Toint, P. L. *Trust-region methods*. (Society for Industrial and Applied Mathematics, 2000).

36 Nicholls, D. G. The influence of respiration and ATP hydrolysis on the proton-electrochemical gradient across the inner membrane of rat-liver mitochondria as determined by ion distribution. *Eur J Biochem* **50**, 305-315, doi:10.1111/j.1432-1033.1974.tb03899.x (1974).

37 Nicholls, D. G. The effective proton conductance of the inner membrane of mitochondria from brown adipose tissue. Dependency on proton electrochemical potential gradient. *Eur J Biochem* **77**, 349-356, doi:10.1111/j.1432-1033.1977.tb11674.x (1977).

38 Nicholls, D. G. Hamster brown-adipose-tissue mitochondria. The control of respiration and the proton electrochemical potential gradient by possible physiological effectors of the proton conductance of the inner membrane. *Eur J Biochem* **49**, 573-583, doi:10.1111/j.1432-1033.1974.tb03861.x (1974).

39 Wang, Z. *et al.* Cyclin B1/Cdk1 coordinates mitochondrial respiration for cell-cycle G2/M progression. *Dev Cell* **29**, 217-232, doi:10.1016/j.devcel.2014.03.012 (2014).

40 Domenech, E. *et al.* AMPK and PFKFB3 mediate glycolysis and survival in response to mitophagy during mitotic arrest. *Nat Cell Biol* **17**, 1304-1316, doi:10.1038/ncb3231 (2015).

41 Harbauer, A. B. *et al.* Mitochondria. Cell cycle-dependent regulation of mitochondrial preprotein translocase. *Science* **346**, 1109-1113, doi:10.1126/science.1261253 (2014).

42 Picard, M., McEwen, B. S., Epel, E. S. & Sandi, C. An energetic view of stress: Focus on mitochondria. *Front Neuroendocrinol* **49**, 72-85, doi:10.1016/j.yfrne.2018.01.001 (2018).

43 Chretien, D. *et al.* Mitochondria are physiologically maintained at close to 50 degrees C. *PLoS Biol* **16**, e2003992, doi:10.1371/journal.pbio.2003992 (2018).

44 Rodenfels, J., Neugebauer, K. M. & Howard, J. Heat Oscillations Driven by the Embryonic Cell Cycle Reveal the Energetic Costs of Signaling. *Dev Cell* **48**, 646-658 e646, doi:10.1016/j.devcel.2018.12.024 (2019).

45 Gibcus, J. H. *et al.* A pathway for mitotic chromosome formation. *Science* **359**, doi:10.1126/science.aao6135 (2018).

46 Cermak, N. *et al.* High-throughput measurement of single-cell growth rates using serial microfluidic mass sensor arrays. *Nat Biotechnol* **34**, 1052-1059, doi:10.1038/nbt.3666 (2016).

47 Olcum, S., Cermak, N., Wasserman, S. C. & Manalis, S. R. High-speed multiple-mode mass-sensing resolves dynamic nanoscale mass distributions. *Nat Commun* **6**, 7070, doi:10.1038/ncomms8070 (2015).

48 Calistri, N. L. *et al.* Microfluidic active loading of single cells enables analysis of complex clinical specimens. *Nat Commun* **9**, 4784, doi:10.1038/s41467-018-07283-x (2018).
